# Specific recognition and ubiquitination of slow-moving ribosomes by human CCR4-NOT

**DOI:** 10.1101/2022.07.24.501325

**Authors:** Eva Absmeier, Viswanathan Chandrasekaran, Francis J O’Reilly, James AW Stowell, Juri Rappsilber, Lori A Passmore

## Abstract

Eukaryotic messenger RNA (mRNA) decay is generally initiated by removal of the 3’ polyadenosine (poly(A)) tail by the CCR4-NOT complex. Yeast Ccr4-Not binds and ubiquitinates ribosomes stalled on mRNAs with sub-optimal codons to trigger deadenylation and decay of the associated transcript. However, the mammalian ortholog of the E3 ubiquitin ligase subunit, CNOT4, is not a constitutive component of human CCR4-NOT. It therefore remains unclear how the mammalian deadenylation machinery targets stalled ribosomes. Here, we reconstitute translational stalling on non-optimal codons. We find that human CCR4-NOT recognizes translating mammalian ribosomes and is required for stable CNOT4 association. Our cryoEM structure reveals that the CNOT3 subunit detects slow translation and locks the L1 stalk of the ribosome in an open conformation to impede further elongation. Using crosslinking mass spectrometry, we show that CNOT4 and CNOT11 also bind in the vicinity of the E site. Overall, our work defines how CCR4-NOT enforces ribosomal stalling in response to low codon optimality.

## Introduction

Protein synthesis is subject to tight regulation to ensure the fidelity of gene expression. mRNA defects (such as truncations, premature stop codons and inappropriate polyadenylation), amino acid starvation, and tRNA insufficiency or poor codon optimality can lead to stalling of ribosomes. This activates quality control pathways to trigger rescue or degradation of ribosomes, and degradation of both the nascent polypeptide chain and the defective mRNA (*1*).

In eukaryotes, mRNAs with sub-optimal codons are subjected to a specialized “codon-optimality decay” pathway (*2, 3*) and codon usage is a major determinant of mRNA half-life (*4–13*). Interestingly, slow translation and thus low codon-optimality may be sensed directly by the conserved multi-functional cytoplasmic deadenylase CCR4-NOT (*14, 15*). Recent cryoEM structures of affinity-purified yeast Ccr4-Not bound to ribosomes revealed that a helical bundle within an N-terminal region (NTR) of the Not5 subunit (CNOT3 in human) directly binds the E site of ribosomes that also contain an empty A site. Not5 binding requires both the E3 ubiquitin ligase subunit, Not4 (CNOT4 in human), and mono-ubiquitination of the 40S eS7 protein (*15*).

The CCR4-NOT complex can be recruited to specific mRNAs to mediate rapid deadenylation and degradation in response to cellular cues (*16*). Its multiple subunits assemble around the large scaffold protein CNOT1 (Fig. 1A) (*17*). The CNOT10 and CNOT11 subunits are absent in yeast (*18*)and bind to an N-terminal region of CNOT1 (*19*), but the exact cellular function of this CNOT10/11 module is unknown. The two poly(A)-specific exonucleases CNOT6/CNOT6L/Ccr4 and CNOT7/CNOT8/Caf1 bind to a central region of CNOT1 to form the nuclease module. CNOT9/Caf40 interacts with CNOT1 downstream of the nuclease module and with a number of RNA-binding proteins to recruit the complex to specific mRNAs. Finally, CNOT2 and CNOT3 form the NOT module with a C-terminal region of CNOT1 (*20*).

**Fig. 1.**
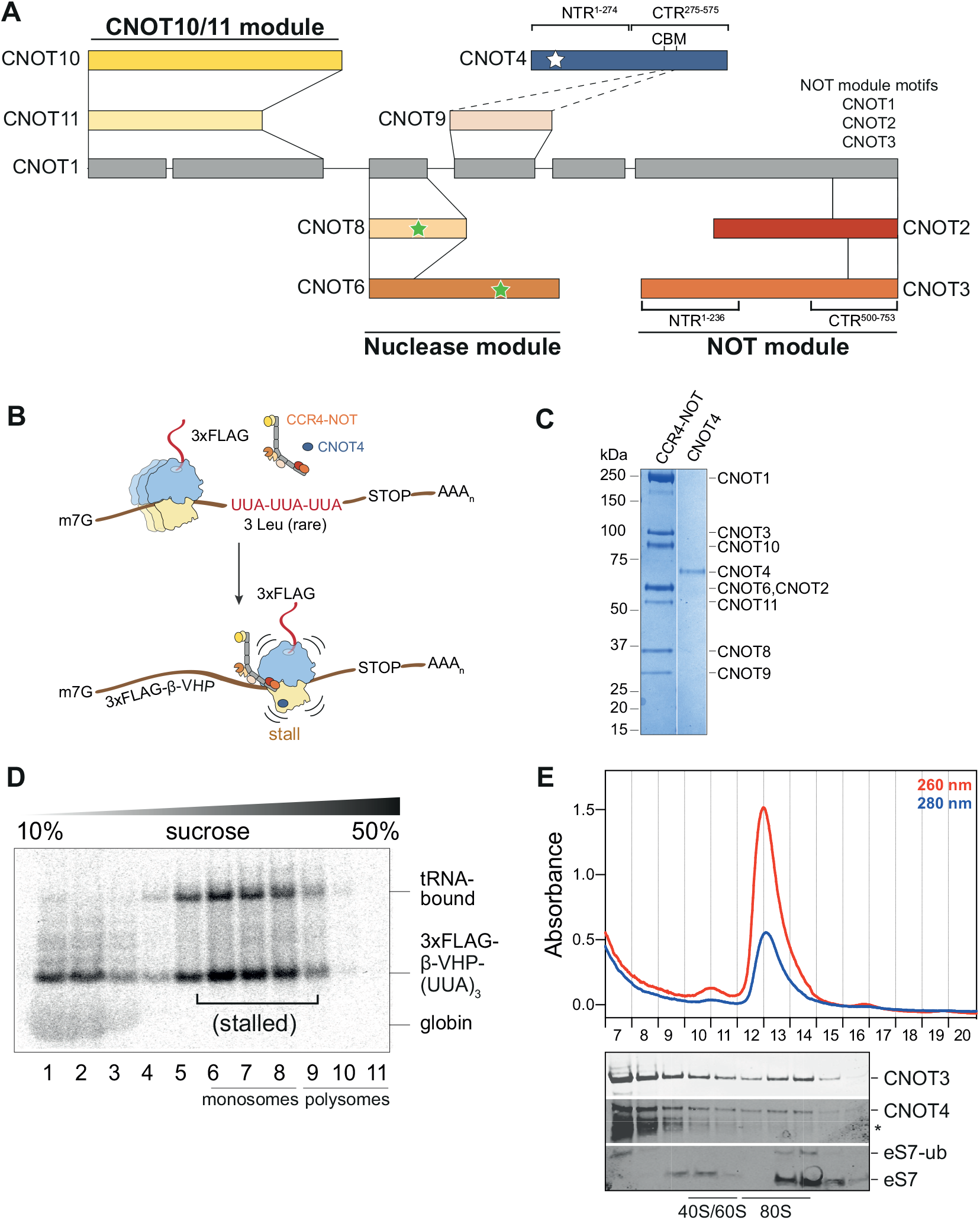
Human CCR4-NOT and CNOT4 recognize stalled ribosomes in an *in vitro* translation system. (**A**) Schematic of subunits and interaction network within human CCR4-NOT. Stars indicate enzymatically active subunits including the E3 ubiquitin ligase and poly(A) specific exonucleases. CBM, Caf40-binding-motif; NTR, N-terminal region; CTR, C-terminal region. (**B**) Schematic of the *in vitro* translation strategy to stall 80S ribosomes on sub-optimal codons in the presence of CCR4-NOT and CNOT4. (**C**) Coomassie-stained SDS-PAGE of purified recombinant CCR4-NOT and its non-constitutive E3 ubiquitin ligase CNOT4. (**D**) Sucrose gradient sedimentation of a translation reaction containing RNCs stalled on *rare* RNA in the presence of 50 nM CCR4-NOT and 75 nM CNOT4. This autoradiogram shows migration of the ^35S^L-Met-labelled nascent polypeptide. (**E**) Polysome profile (Absorbance 260 nm red, 280 nm blue) of fractions 7-20 (top) and Western immunoblots of fractions 7-16 probed with antibodies against CNOT3, CNOT4 and the ribosomal protein eS7 (bottom). An asterisk denotes a CNOT4 antibody cross-reacting band.

Unlike yeast Not4, the human E3 ubiquitin ligase CNOT4 is not an integral subunit of CCR4-NOT (*18, 21, 22*). Interestingly, deletion of the CNOT4 NTR results in stable binding of the C-terminal region (CTR) to CCR4-NOT, indicating that the CNOT4 NTR has an auto-inhibitory role (*23*). In metazoans, CNOT4 has several binding interfaces located in its CTR, including a conserved sequence motif (the Caf40-binding-motif or CBM) that directly interacts with CNOT9 (*23*) (Fig. 1A). Overall, the core of yeast and human CCR4-NOT complexes is conserved but there are differences in their subunit composition and arrangement.

Given the differences between yeast and human CCR4-NOT, it is unknown how the human complex mediates codon-optimality decay. Moreover, it remains unclear how CCR4-NOT, CNOT4 and other translational control factors exhibit selectivity, namely the ability to spare actively translating ribosomes and specifically target only the minor population of ribosome-nascent chain complexes (RNCs) that require rescue and/or mRNA decay. Here, we establish an *in vitro* reconstitution approach to permit mechanistic dissection of the interplay between mammalian CCR4-NOT and translation. We reveal the molecular features of slow-moving mammalian 80S ribosomes that serve as substrates for CCR4-NOT and CNOT4. We also show how CCR4-NOT enforces stalling and determine the role of ubiquitination in CCR4-NOT binding.

## Results & Discussion

### Human CCR4-NOT and CNOT4 bind and ubiquitinate stalled mammalian 80S ribosomes

To gain mechanistic insight into the link between CCR4-NOT, CNOT4 and codon-optimality, we reconstituted translational stalling on sub-optimal codons in rabbit reticulocyte lysate (RRL) (Fig. 1B). We exploited the fact that the RRL is a minimal system that has downregulated the quantities of most tRNAs except those necessary to decode the efficiently translated α- and β-globin mRNAs. RRL pre-treated with nuclease to degrade the globin transcripts can be employed to investigate an mRNA of choice (*24*). To study codon-optimality decay, we used a short mRNA encoding the cytosolic domain of Sec61b with a stretch of three consecutive UUA leucine codons before the stop codon, which we termed *‘rare’* mRNA (Fig. 1B). The UUA codons are decoded by tRNA^Leu,UAA^, which is poorly abundant in the RRL (*24*). To facilitate affinity purification of RNCs, the model mRNA encoded an N-terminal 3x FLAG tag and a stably folded villin head piece (VHP) domain (*25*). The mRNA was capped with a 5’-cap analogue and polyadenylated using *Escherichia coli* poly(A) polymerase to contain a 3’-poly(A) tail.

We performed *in vitro* translations using the *rare* mRNA in the presence of ^35S^L-methionine, which is incorporated into the nascent chain during translation. We subsequently separated the reactions on sucrose gradients and analyzed the fractions by autoradiography. We found that translation of the *rare* mRNA in RRL resulted in stalling at the UUA triplet locus and an accumulation of RNCs in the deeper fractions of the gradient (Fig. S1A). Translational stalling was not observed when the RRL was supplemented with total tRNAs purified from pig liver, which contains tRNA^Leu,UAA^ in high amounts (Fig. S1B). Thus, tRNA insufficiency mimics low codon-optimality and stalls the ribosome, which remains bound to the nascent polypeptide chain.

To test whether CCR4-NOT and CNOT4 target the stalled ribosomes, we purified recombinant human CCR4-NOT produced using baculovirus-mediated overexpression in insect cells (*26*) and recombinant human CNOT4 expressed in *Escherichia coli* (Methods; Fig. 1C). We then performed *in vitro* translation reactions in the presence of CCR4-NOT and CNOT4. We estimated the cellular concentrations of CCR4-NOT to be on the order of 20 nM (*27*) and therefore used a slight excess of 50 nM CCR4-NOT and 75 nM CNOT4 in our assays. As controls to test whether CCR4-NOT and CNOT4 specifically target stalled ribosomes we used globin mRNAs from non-nuclease treated RRLs, which are translated very efficiently.

First, we found that addition of 50 nM CCR4-NOT and 75 nM CNOT4 to the translation reaction did not appear to alter translation and stalling of RNCs on the *rare* mRNA (Fig. 1D, compare to Fig. S1A). Next, we analyzed the migration of CCR4-NOT and CNOT4, and monitored eS7 ubiquitination by CNOT4 using sucrose gradients and Western immunoblotting. Both CCR4-NOT and CNOT4 co-migrate with 80S ribosomes on *rare* mRNA (Fig. 1E and S1C). A small fraction of the eS7 ribosomal protein was mono-ubiquitinated, indicating that CNOT4 ubiquitinates stalled RNCs, but not the idle 80S ribosomes that predominate the RRL.

Given that CNOT4 is not a constitutive subunit of human CCR4-NOT, we also tested whether CCR4-NOT is required for CNOT4 binding and eS7 ubiquitination and *vice versa*. CNOT3 co-migrates with *rare* mRNA-bound RNCs, even when CNOT4 and eS7 ubiquitination are absent (Fig. S1D). In contrast, yeast Not4 was required for stable association of Not5 with stalled ribosomes in a previous study (*15*). This suggests that human CCR4-NOT binds more stably to 80S ribosomes. CNOT4 promotes eS7 mono-ubiquitination independent of CCR4-NOT, but stable association of CNOT4 to RNCs requires CCR4-NOT (Fig. 1E and S1E). When we analyzed the globin mRNA, CCR4-NOT and CNOT4 also bound to RNCs and eS7 ubiquitination occurred (Fig. S1F-H). However, both were much less efficient compared to *rare* mRNA (compare CNOT3 and CNOT4 levels in 80S fractions of Fig. S1C and S1H), supporting the idea that CCR4-NOT and CNOT4 specifically target stalled RNCs.

Together, our data suggest that human CCR4-NOT recognizes and co-migrates with stalled 80S ribosomes, and that CNOT4 is required for eS7 ubiquitination in the RRL.

### Human CCR4-NOT recognizes stalled ribosomes with the CNOT3 NTR

To determine the molecular basis for how human CCR4-NOT recognizes and ubiquitinates 80S ribosomes that have stalled on sub-optimal codons, we employed single particle cryoEM analyses of purified RNC complexes. Ribosomes translating the *rare* mRNA in RRL were pelleted through a sucrose cushion and RNCs were enriched using FLAG affinity pulldowns on the nascent chain and eluted using 3xFLAG peptide. A subset of these ribosomes is expected to be stalled, eS7-ubiquitinated and bound by CCR4-NOT and CNOT4. Samples for cryoEM were stabilized by bis(sulfosuccinimidyl)suberate (BS3) crosslinking during pelleting and during pellet resuspension. After processing the cryoEM data by 2D and 3D classification, focused classification with signal subtraction around the P site, and subsequently, on the E site, we obtained a cryoEM map of ribosomes bound to CCR4-NOT at 3.1 Å resolution (Fig. S2, S3).

The structure revealed that the stalled ribosome is in a canonical state with the peptidyl tRNA^Leu,UUA^ in the P/P state and with the A and E sites devoid of tRNAs (Fig. 2A). Based on the results we obtained from our *in vitro* translation experiments and in agreement with our experimental design (stalling on the UUA codons of the *rare* mRNA), we modeled a nascent chain with 2 leucyl residues and a UUA codon (Fig. S4A-B).

**Fig. 2.**
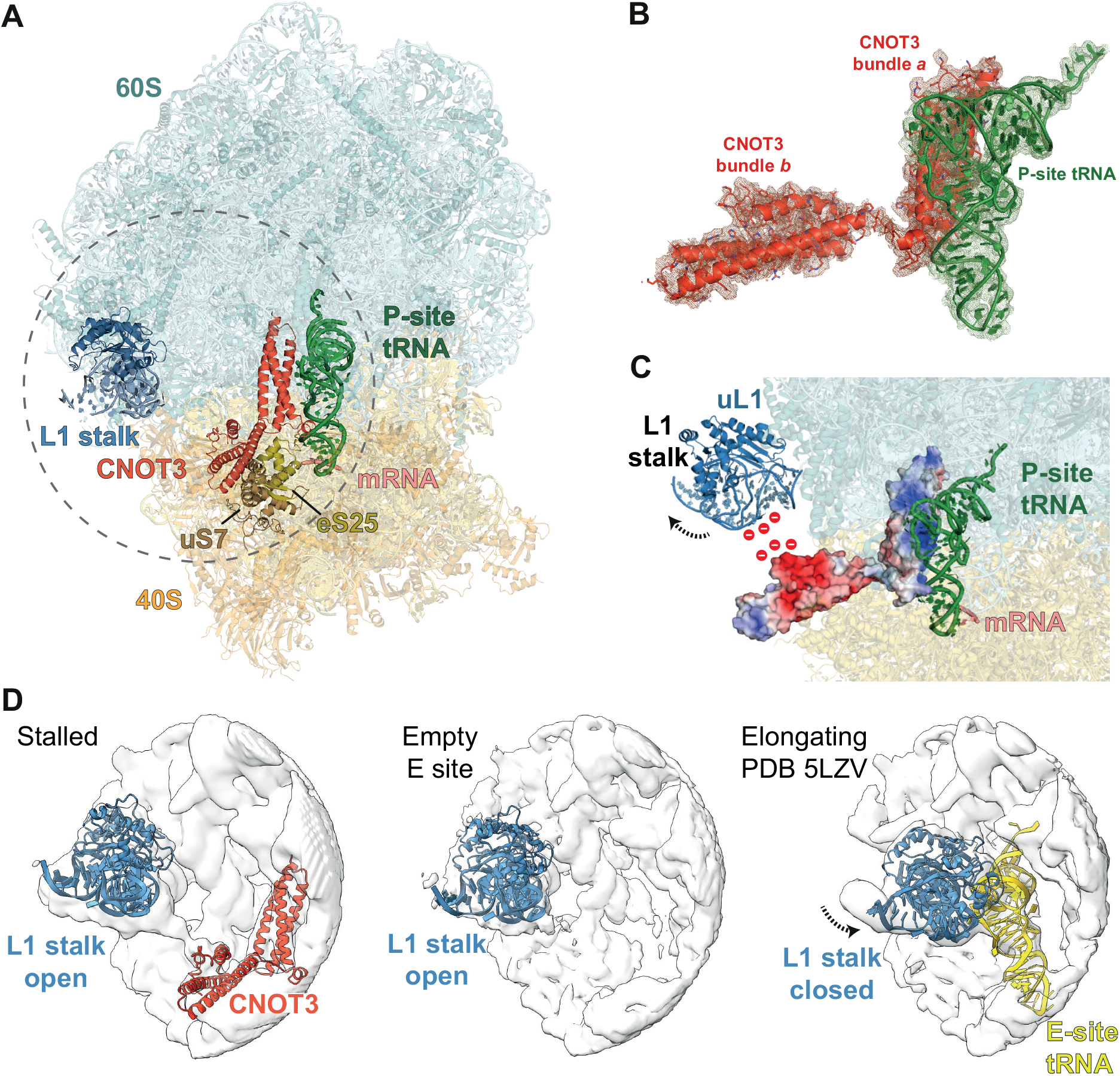
Structure of mammalian CCR4-NOT-bound to the 80S ribosome. (**A**) Overview of the atomic model of stalled 80S ribosomes containing the CNOT3 subunit of CCR4-NOT in the E-site, obtained using cryoEM. 60S, cyan; 40S, yellow; CNOT3, orange; L1 stalk of the ribosome, blue; P-site tRNA, green; mRNA, salmon; eS25, gold; uS7, olive. (**B**) Atomic model of CNOT3 and P-site tRNA shown in the cryoEM map. Helical bundles *a* and *b* are labelled. Map was calculated from 2-fold downscaled particles (to a pixel size of 1.66 Å/pix) to improve interpretability and is contoured at 5 r.m.s.d. (**C**) The surface charge of CNOT3 mapped onto its structure. A conserved negatively-charged patch on CNOT3 repels and holds the L1 stalk open (red, negative charge; blue positive charge). (**D**) Comparison of the L1 stalk positions in surface outline representation between CNOT3-bound stalled ribosomes (left), stalled ribosomes from the same dataset but with empty E site (middle) and ribosomes stalled during elongation at the stop codon (PDB 5LZV; right).

The N-terminal region (NTR) of CNOT3 (residues 1-236) binds the vacant E site of the 80S ribosome (Fig. 2A-B). An N-terminal 3-helix bundle (bundle *a*) of CNOT3 (residues 1-111) packs against the P-site tRNA and makes additional contacts to the 28S rRNA via a conserved positively-charged surface patch (Fig. 2B-C, S4C-D). Comparison of bundle *a* with Not5 bundle *a* from the yeast ribosome structure shows that they are structurally conserved (0.6 Å r.m.s.d), consistent with a sequence identity of 54.5% (Fig. S5A-B). A second domain of CNOT3, which was not well-ordered in the yeast structure, could be modelled as an additional three-helix bundle (bundle *b*) (residues 115-236) oriented at an angle of 105° to bundle *a*. Helical bundle *b* is less conserved than bundle *a* and shares a sequence identity of 29.8% with yeast. Bundle *b* interacts with uS7 and eS25 on the head of the 40S subunit (Fig. 2A).

Interestingly, a highly conserved negatively-charged patch on helical bundle *b* (Fig. 2C and S5C) is located next to the L1 stalk. We propose that this repels the negatively-charged phosphate backbone of the rRNA in the L1 stalk and holds the latter in a fully open conformation. This L1 stalk conformation is reminiscent of stalled ribosomes with an empty E site, such as during translation of poly(A) stretches (*28*). The open L1 stalk conformation does not persist stably during normal elongation (Fig. 2D) and therefore is likely stabilized by the negatively-charged surface on CNOT3, potentially to lock ribosomes in this conformation while awaiting downstream steps. The corresponding helical bundle *b* in yeast is approximately 20 residues shorter and an Alphafold2 (*29, 30*) prediction reveals that this results in shortening of two the long helices (Fig. S5A-B). Comparing the electrostatic surface charge of yeast and human bundle *b* reveals that the overall negative surface charge of bundle *b* is conserved, but that human bundle *b* has a positive surface patch on the tip of the two long helices that is absent in yeast (Fig. S5C-D). Additional contacts between the extended CNOT3 helices and the C-terminus of eS25 may stabilize this position of helical bundle *b* and the L1 stalk in human relative to yeast. These additional contacts would be consistent with the higher affinity of human CCR4-NOT for 80S ribosomes.

Interestingly, the CNOT3 NTR is a hotspot for disease mutations (Fig. S5A). Residues E20 and L48 in bundle *a*, and K119, E147 and R188 in bundle *b* have been found to be mutated in developmental disorders (*31*). Residues E20, R57 and E70 in bundle *a* are frequently mutated in somatic cancer (*32*). This underlies the importance of this region for CNOT3 function and suggests that disruption of the ribosome-interacting region results in physiological defects.

Together, our structure of human CCR4-NOT bound to stalled ribosomes reveals that the CNOT3 NTR is highly conserved and has evolved to detect and stably bind ribosomes stalled during elongation in the canonical state and with vacant A and E sites.

### CCR4-NOT and CNOT4 are both positioned around the E site

The cryoEM structure provides important insight into the molecular recognition of stalled 80S ribosomes by the NTR of CNOT3. However, the other seven subunits of CCR4-NOT and CNOT4 were not visible in the cryoEM maps. Presumably, they are either unstable during cryoEM specimen preparation or they engage in a flexible manner with the 80S and are blurred during single particle averaging. To determine whether other CCR4-NOT subunits and CNOT4 make specific interactions with 80S ribosome in solution, we performed crosslinking mass spectrometry analyses of CCR4-NOT-, and CNOT4-bound RNCs.

Several CCR4-NOT subunits, including CNOT3, CNOT1, CNOT10, and CNOT11, as well as CNOT4, crosslinked to stalled 80S ribosomes (Fig. 3 and S6). The CNOT3 NTR crosslinked in the vicinity of the E site, the L1 stalk and the interface between the small and large subunits, including to ribosomal proteins uS13, uL1, eS25, uL5, eL42, eL29 and uL11, which agrees with the density we observed for CNOT3 in cryoEM (Fig. 3A and S6A). CNOT4 crosslinked to 40S proteins eS28 and eS1, located near its substrate eS7 and the CNOT3 NTR. Intriguingly, the NTR of CNOT4, which contains the RING E3 ligase domain, but not the CTR, crosslinked to the ribosome (Fig. 3B and S6A).

**Fig. 3.**
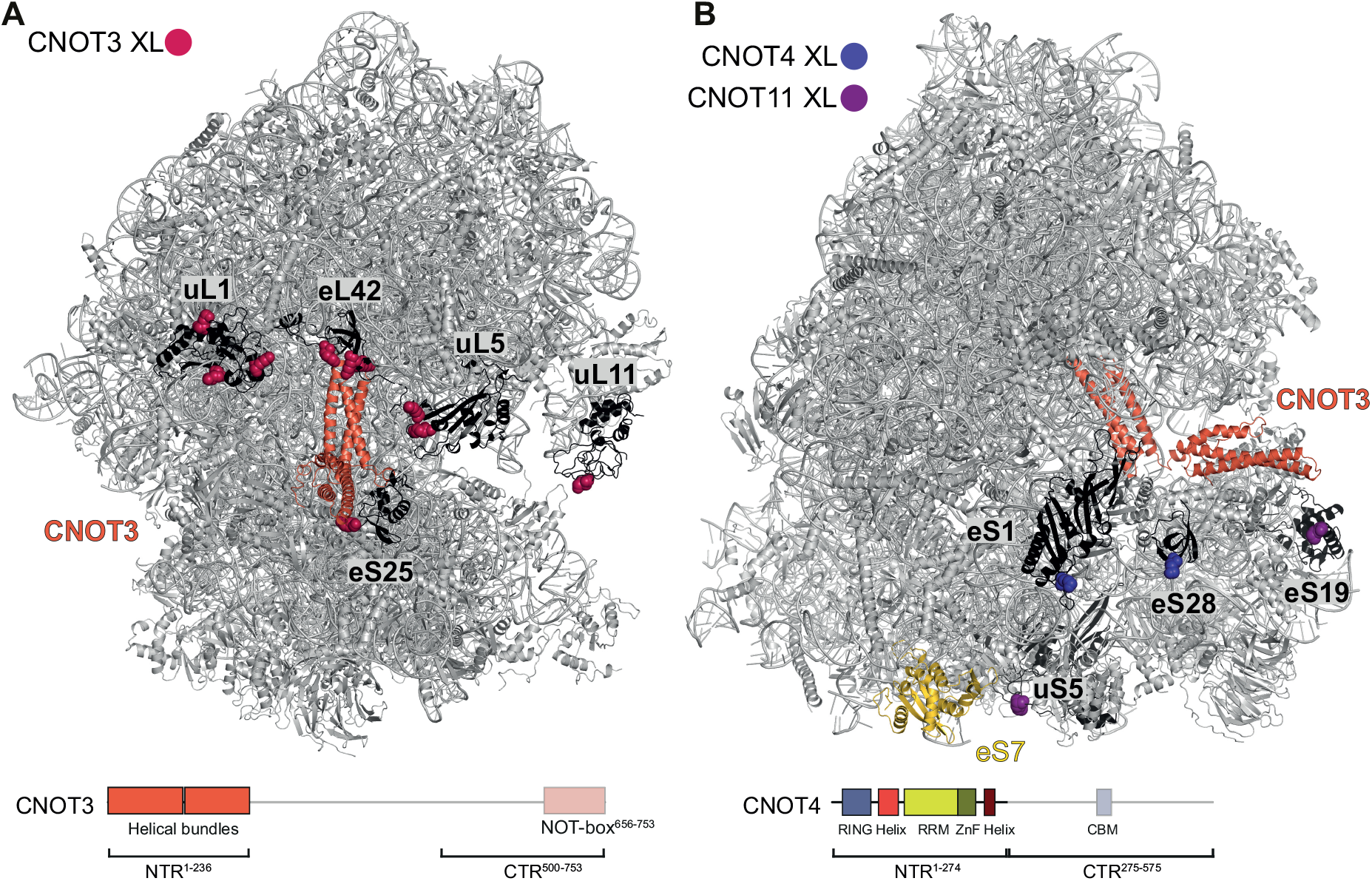
CCR4-NOT binds 80S ribosomes in the vicinity of the E site. Ribosomal proteins that crosslink to (**A**) CNOT3, (**B**) CNOT4 and CNOT11 are colored black within the structure of the 80S ribosome (grey) bound to CNOT3 (orange) (top). The ubiquitination substrate of CNOT4, eS7, is colored gold in panel (**B**). Crosslinked residues within these proteins are represented as spheres and are colored according to the CCR4-NOT subunit as indicated. Domain diagrams of CNOT3 (**A**) and CNOT4 (**B**) are shown at the bottom. Regions that crosslink to the 80S are shown opaque. Protein residues of eL29, uS13 and eS31, which crosslinked to CNOT3 and the CNOT10/11 module, were not modelled and therefore are not mapped onto the structure. RRM, RNA recognition motif; ZnF, zinc finger; CBM, Caf40-binding motif; NTR, N-terminal region; CTR, C-terminal region.

The CNOT10/11 module also crosslinked to stalled ribosomes, specifically to uS5, eS19, eS31and eL29 (Fig. 3B and S6A). Thus, the CNOT10/CNOT11 module resides in close proximity to the 40S subunit. Intriguingly the CNOT1 N-terminus crosslinked to ubiquitin, which is presumably attached to eS7.

The crosslinking data also provided information on the arrangement of CCR4-NOT subunits and CNOT4 when bound to stalled ribosomes (Fig. S6B-C). First, the CNOT3 NTR crosslinked to the CNOT10/11 module and the CNOT4 NTR. This is in agreement with these proteins contacting the same region of the 80S ribosome and therefore being located in close proximity to each other on stalled ribosomes (Fig. 3). The CNOT3 NTR also crosslinked to CNOT2, CNOT1 and the exonuclease subunit CNOT6. The C-terminal region of CNOT3 crosslinked to a number of subunits including CNOT2 and CNOT1 (Fig. S6B-C), in agreement with a previously-reported crystal structure of the NOT module (*33*). The CNOT10/11 module is located in close proximity to the NOT module. The CNOT3 CTR also crosslinked to CNOT4 and CNOT9.

Taken together these data suggest that the region around the E site and the 40S subunit form an interaction hotspot for CCR4-NOT, especially to CNOT3, CNOT4 and the CNOT10/11 module. The crosslinking mass spectrometry data suggest that CNOT4 and CNOT3 directly interact with each other. These contacts may be important for the requirement of CCR4-NOT to promote stable CNOT4 binding to stalled ribosomes.

### Human CNOT4 interacts with both CCR4-NOT and the 80S ribosome

Our crosslinking mass spectrometry data suggest that the CNOT4 NTR binds to the ribosome directly, in the close vicinity of eS7 and the E site. In addition to the N-terminal RING, the CNOT4 NTR (residues 1-274) contains a long α-helix, an RNA recognition motif (RRM) domain and a zinc-finger (ZnF) (Fig. 4A). The precise function of the RRM and the ZnF are not known but they are predicted to form a single structural unit (Fig. 4B). The predicted structure of the CNOT4 NTR has a large positively-charged surface encompassing regions of the linker, the RRM and the ZnF, which may play a role in ribosome recognition. The CNOT4 CTR (residues 275-575) is predicted to be mainly disordered, except for an α-helix (Caf40-binding motif, CBM), which is known to bind to CNOT9 (*23*).

**Fig. 4.**
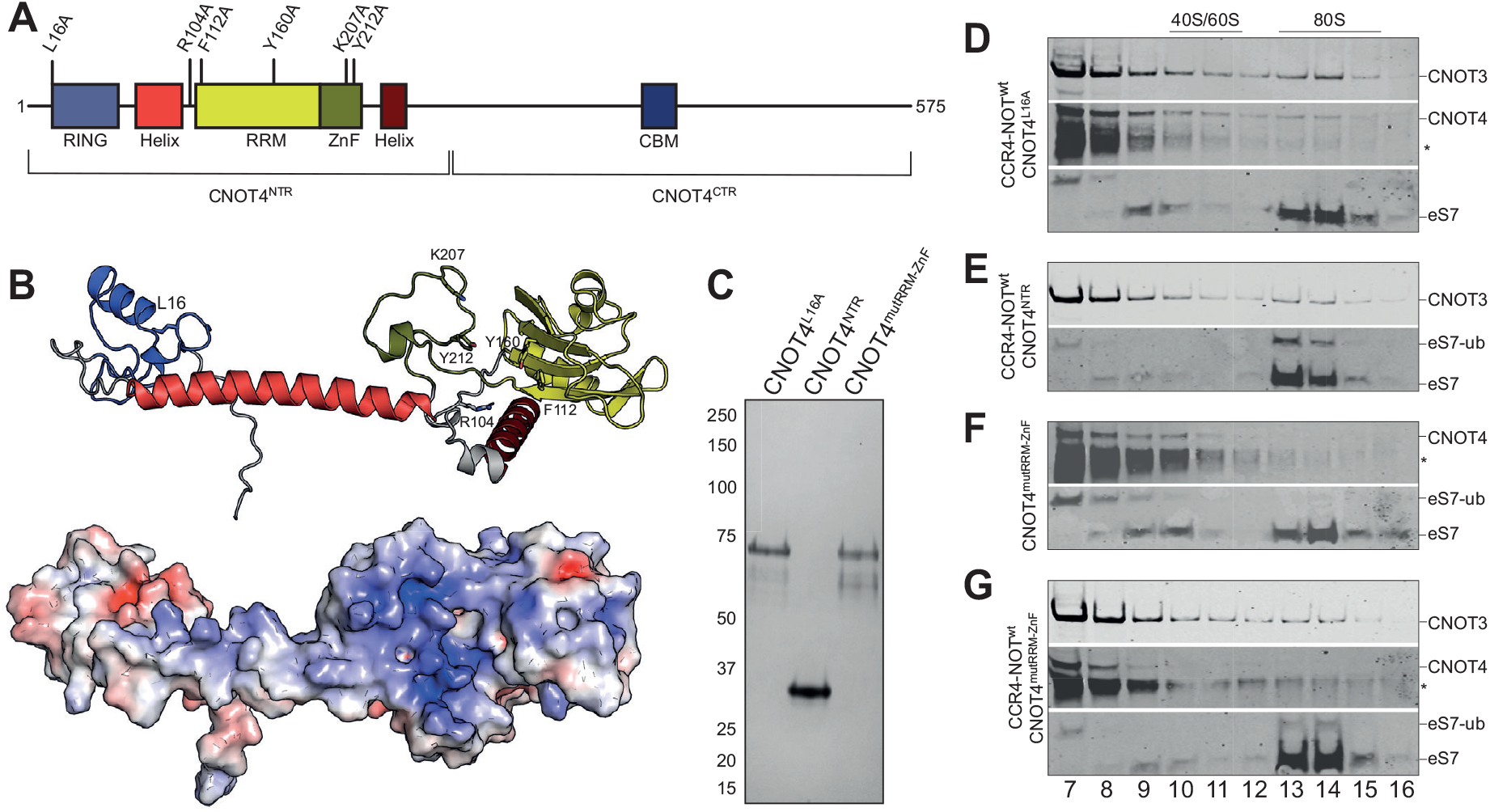
The CNOT4 RRM and ZnF domains are required for association with the 80S ribosome and eS7 ubiquitination. (**A**) Human CNOT4 domain arrangement and positions of mutations. RING, really interesting new gene; RRM, RNA recognition motif; ZnF, zinc finger; CNOT4^NTR^, N-terminal region (1-274); CNOT4^CTR^, C-terminal region (275-575). Mutated residues are indicated. (**B**) Alphafold2 prediction of human CNOT4^NTR^, colors as in panel (A) (top) and electrostatic surface potential of CNOT4^NTR^ (blue, positive; red, negative). (**C**) Representative Coomassie-stained SDS-PAGE of purified CNOT4 variants: CNOT4^L16A^, CNOT4^NTR^, CNOT4^mutRRM-ZnF^ (R104A, F112A, Y160A, K207A, Y212A). (**D-G**) Western immunoblots of *in vitro* translation reactions with *rare* mRNA, 50 nM CCR4-NOT and 75 nM CNOT4 variants, resolved by sucrose gradient sedimentation (fractions 7-16). (**D**) CCR4-NOT and CNOT4^L16A^, (**E**) CCR4-NOT and CNOT4^NTR^, (**F**) CNOT4^mutRRM-ZnF^, (**G**) CCR4-NOT and CNOT4^mutRRM-ZnF^ Fractions were probed using antibodies against CNOT3, CNOT4 and eS7. Asterisks denote bands that cross-react with the CNOT4 antibody.

To investigate CNOT4 binding and ubiquitination of the ribosome, we designed several CNOT4 mutants. First, to determine whether CNOT4-mediated ubiquitination is required for CNOT4 association with CCR4-NOT-RNC complexes in higher eukaryotes, we designed a CNOT4 mutant (CNOT4^L16A^) that cannot recruit the cognate E2 ubiquitin conjugating enzyme (*34*). Second, we produced a CNOT4 variant comprising only the NTR (CNOT4^NTR^) to test whether this minimal construct can ubiquitinate eS7. Finally, we introduced mutations in positively-charged and aromatic residues (R104A in the linker, F112A and Y160A in the RRM, K207A and Y212A in the ZnF) on the surface of CNOT4 (CNOT4^mutRRM-ZnF^) (Fig. 4A-B). We then tested whether purified CNOT4 variants (Fig. 4C) bind the ribosome and whether they promote ubiquitination of eS7 (Fig. 4D-G, S7).

In the presence of CCR4-NOT, CNOT4^L16A^ co-migrates with the 80S ribosome, but does not ubiquitinate eS7 (Fig. 4D), suggesting that, unlike in yeast (*15*), eS7 ubiquitination is not required for CCR4-NOT and CNOT4 association with stalled ribosomes. CNOT4^NTR^ efficiently ubiquitinates eS7 (Fig. 4E) but we were unable to monitor its association with ribosome owing to a lack of a suitable antibody.

Despite carrying an intact RING domain, CNOT4^mutRRM-ZnF^ did not ubiquitinate eS7 in the absence of CCR4-NOT, and did not co-migrate with 80S ribosomes (Fig. 4F). In contrast, wild-type CNOT4 ubiquitinated eS7 in the absence of CCR4-NOT (Fig. S1E). Intriguingly, when CCR4-NOT was included in the translation reaction, CNOT4^mutRRM-ZnF^ ubiquitinated eS7 (Fig. 4G). However, we did not observe any CNOT4 ^mutRRM-ZnF^ in the 80S fractions.

Together, these results indicate that stable binding of CNOT4 to 80S ribosomes seems to require two interaction sites. Firstly, because eS7 is ubiquitinated by wild-type but not CNOT4 ^mutRRM-ZnF^ in the absence of CCR4-NOT, the CNOT4 NTR likely interacts with the ribosome through charged and aromatic residues on the surface of the RRM and the ZnF. This agrees with the observed crosslinks between CNOT4 and the ribosome. Secondly, because eS7 ubiquitination by CNOT4 ^mutRRM-ZnF^ is rescued in the presence of CCR4-NOT, the CNOT4 CTR likely interacts with CCR4-NOT-bound ribosomes, possibly through direct interactions with CNOT3 (Fig. S6B-C). CNOT4 NTR and CTR interactions are required for stable CNOT4 interaction with the ribosome.

### Model for recognition of stalled ribosomes by human CCR4-NOT and CNOT4

Here, we established an *in vitro* system to trap human CCR4-NOT and the non-constitutive E3 ubiquitin ligase CNOT4 subunit during specific recognition and ubiquitination of mammalian ribosomes stalled on sub-optimal codon stretches. Thus, our work identifies human CCR4-NOT and CNOT4 as factors crucial for the codon-optimality decay pathway in mammals.

Our data reveal that codon-optimality decay is generally conserved, but there are also striking differences between yeast and mammals. This includes the presence of the metazoan-specific CNOT10 and CNOT11 subunits. In addition, Not4 is a constitutive subunit of yeast Ccr4-Not but not human CCR4-NOT, and eS7 ubiquitination is required for stable association of the yeast, but not human, complex. Together, mammalian CCR4-NOT and CNOT4 have evolved to form a modular machinery that directs complex gene expression regulation in higher eukaryotes. We propose the following model for mammalian CCR4-NOT and CNOT4 as codon-optimality decay factors in mammals.

Elongating ribosomes do not persist with empty A sites when translating optimal codon stretches owing to efficient decoding of the A-site codon by the abundant eEF1A·aminoacyl-tRNA·GTP ternary complex. This allows decoding to proceed in a timely manner, followed by accommodation, peptidyl transfer, subunit rotation and eEF2·GTP-mediated translocation to the next codon (Fig. 5, top). Slower elongation on sub-optimal codon stretches (such as the UUA triplet used here) increases the time that ribosomes spend in the post-translocation state with an empty A site, while they await the cognate aminoacyl tRNA. If a cognate aminoacyl tRNA does not arrive (e.g. in the case of low codon optimality or tRNA insufficiency), the L1 stalk opens and the E-site tRNA dissociates (Fig. 5, bottom).

**Fig. 5.**
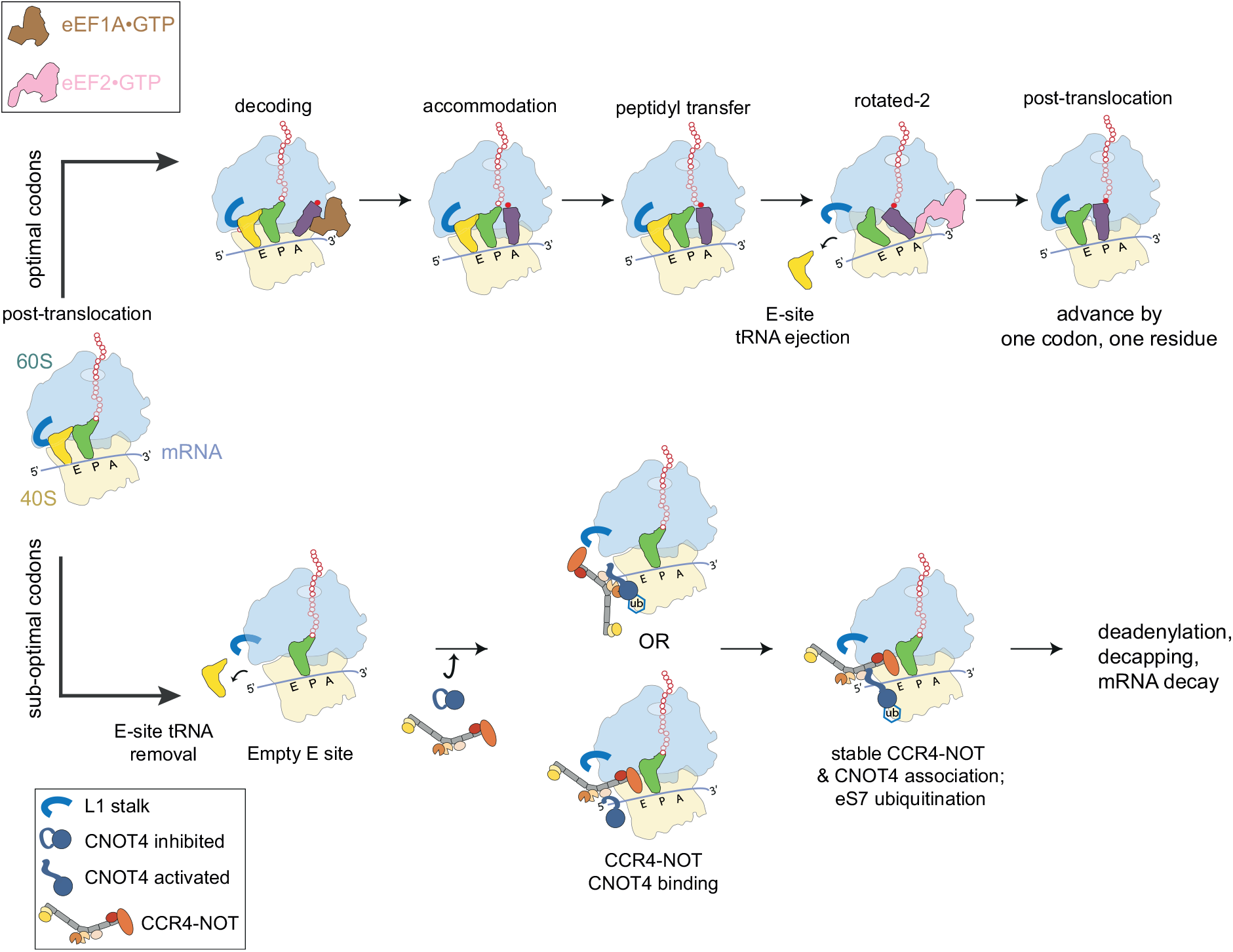
Model for recognition of slow-moving ribosomes by human CCR4-NOT and CNOT4. Elongating ribosomes in the canonical state with peptidyl-tRNA (green) in the P site and deacylated tRNA (gold) in the E site await the next cognate aminoacyl-tRNA (purple) (top, left). Various aminoacyl-tRNAoeEF1AoGTP ternary complexes sample the A site codon during decoding until the cognate aminoacyl-tRNA arrives and accommodates upon GTP hydrolysis and egress of eEF1A (top). Peptidyl transfer to lengthen the nascent chain (hollow red beads) by one residue (solid red bead), subunit rotation and formation of hybrid state A/P and P/E tRNAs occur spontaneously. eEF2oGTP is recruited to this state and uses GTP hydrolysis to mediate translocation by advancing the mRNA by one codon to reset the cycle to the next post-translocation state. When elongating through stretches of sub-optimal codons (bottom), the post-translocation state persists, and the E-site tRNA leaves. CCR4-NOT and CNOT4 can bind and mono-ubiquitinate eS7. It remains unclear whether CNOT4 recruits CCR4-NOT or *vice versa*. This triggers poly(A) tail removal, decapping and mRNA decay by exonucleases.

The conserved CNOT3 NTR is a highly specialized molecular sensor and effector, which recognizes the empty E site of slow-elongating ribosomes *via* a conserved positive surface of helical bundle *a*. We show that CNOT3 stabilizes the L1 stalk in an open conformation that stalls further elongation *via* a conserved negatively-charged surface on helical bundle *b*. The CNOT10/11 module may also engage in interactions with 40S subunit proteins. CNOT10 and CNOT11 orthologues can be found in all eukaryotes, except in fungi, and thus far no function has been assigned to these proteins. Our data suggest that they may act as additional adaptors to the ribosome and they may recruit additional downstream factors.

The CNOT4 NTR (specifically the RRM and ZnF domains) interacts with the 40S in close proximity to the CNOT3 NTR. It is possible that CNOT4 binds stalled ribosomes prior to CCR4-NOT *via* its NTR, which would relieve CNOT4 auto-inhibition, recruit CCR4-NOT and allow formation of a stable CCR4-NOT·CNOT4·80S complex (Fig. 5, bottom). In agreement with this, CNOT4 mono-ubiquitinates eS7, even in the absence of CCR4-NOT. However, unlike in yeast, eS7 ubiquitination is not required for CNOT4 and CCR4-NOT comigration with RNCs, as shown with the CNOT4^L16A^ variant.

Alternatively, CCR4-NOT may recognize stalled ribosomes through the CNOT3 NTR prior to CNOT4 binding. CNOT4 would then sense CNOT3 in the E site, which together with the 80S ribosome, may form a composite binding site, allowing CNOT4 to dock onto the stalled ribosome. Auto-inhibition would be alleviated and CNOT4 could ubiquitinate eS7 (Fig. 5, bottom). In agreement with this, CCR4-NOT binds stalled ribosomes in the absence of CNOT4 and is required for stable CNOT4 binding.

Together, binding of CCR4-NOT and CNOT4, and eS7 ubiquitination likely commit the bound transcript to mRNA decay via deadenylation, followed by 5’ decapping and degradation. Our reconstituted system sets the stage to address the roles of other CCR4-NOT subunits and the downstream steps in codon-optimality decay.

## Materials and Methods

### Protein production

#### Human CNOT4

Synthetic human *CNOT4*, optimized for expression in *Escherichia coli*, was cloned into a FX cloning vector (gift from Dr. Harvey MacMahon, MRC-LMB) under the control of a T7 promotor to produce recombinant protein containing a SUMO-cleavable N-terminal His6-SUMO tag. Amino acid mutations were generated with QuickChange mutagenesis, *CNOT4* truncations were PCR amplified and cloned into a FX vector.

Proteins were expressed in *Escherichia coli* BL21 (DE3) in 6 litres Terrific Broth (TB) medium. Cells were grown to an OD_600_ of 0.7 at 37 °C, briefly cooled down and then induced with 1 mM IPTG and grown over night at 18 °C. Cells were harvested and frozen in liquid nitrogen until further use.

All purification steps were performed at 8 °C and proteins were kept on ice. Cell pellets were thawed and resuspended in 200 ml buffer A (200 mM NaCl, 40 mM HEPES-NaOH pH 7.5, 20 mM imidazole, 2 mM magnesium acetate, 0.1 mM TCEP), supplemented with EDTA-free protease inhibitors (Roche) and 2 mg DNaseI, and lysed by sonication. The lysate was cleared by 30 min centrifugation at 21,000 rpm. The cleared lysate was incubated with 10 ml of a 50% Ni-NTA solution on a tube roller for 1 h. The lysate-Ni-NTA mix was separated via a gravity flow column. The beads were washed with 5 x 25 ml buffer A. The sample was eluted with 5 x 10 ml buffer B (200 mM NaCl, 40 mM HEPES-NaOH pH 7.5, 250 mM imidazole, 2 mM magnesium acetate, 0.1 mM TCEP). SUMO protease was added at a 1:30 ratio to the elution and the tag was cleaved over-night on ice.

The cleaved protein was subsequently purified via a 5 ml ButylFF column preequilibrated in buffer C (600 mM ammonium sulfate, 40 mM HEPES-NaOH pH 7.5, 0.1 mM TCEP). Prior to loading, the protein was supplemented with 600 mM ammonium sulfate. The protein was eluted over a linear 12 column volume (CV) gradient to 100% buffer D (40 mM HEPES-NaOH pH 7.5, 0.1 mM TCEP). The protein-containing fractions were pooled, diluted to 80 mM salt and loaded onto a 5 ml Heparin HP column pre-equilibrated in buffer E (80 mM NaCl, 40 mM HEPES-NaOH pH 7.5, 2 mM magnesium acetate, 0.1 mM TCEP). The protein was eluted with a linear gradient over 12 CV to 100% buffer F (1 M NaCl, 40 mM HEPES-NaOH pH 7.5, 2 mM magnesium acetate, 0.1 mM TCEP). The protein was concentrated and run on a S75 16/60 gel filtration column, equilibrated in gel filtration buffer (200 mM NaCl, 20 mM HEPES-NaOH pH 7.5, 0.1 mM TCEP). The protein-containing fractions were pooled, concentrated in a 30 kDa MW cut-off centrifugation filter, flash frozen in liquid nitrogen and stored at −80°C until further usage.

#### Human CCR4-NOT complex

Synthetic genes encoding human CCR4-NOT proteins (*CNOT1, CNOT2, CNOT3, CNOT6, CNOT8-SII, CNOT9, CNOT10, CNOT11*), optimized for expression in *E. coli*, were cloned under the control of a polH promotor into a pACEBac vector and subsequently combined for co-expression into a single modified bigBac vector via Gibson assembly(*26*). *CNOT8* also encoded a C-terminal PreScission cleavable StrepII-tag.

V0 virus was produced by transfecting 1.5 ml of Sf9 cells at 0.7×10^6^ cells/ml with 7 μg of plasmid, mixed with 13 μl Fugene and 100 μl SF900 medium in one well of a 6-well plate. For one expression, a minimum of three wells was transfected. Cells were topped up the next day with 1.5 ml fresh SF900 medium and harvested 60 h post-transfection. Human CCR4-NOT complex was then produced using a low MOI (multiplicity of infection) strategy: 150 ml of Sf9 cells at 1.5×10^6^ cells/ml were infected with 1 ml of fresh V0 virus. Cells were diluted after 24 h with 150 ml, and subsequently at 48 h with 200 ml SF900 medium, to maintain a cell density between 1.5-3 x10^6^ cells/ml. Cells were harvested 72-84 h postinfection at a viability of 92-95% at 800*g* for 10 min, resuspended in ice cold PBS, pelleted at 800*g* and then flash frozen for storage at −80 °C. For one CCR4-NOT purification 6 litres of expression culture was used.

For protein purification, cell pellets were resuspended in 250 ml lysis buffer (150 mM NaCl, 50 mM HEPES-NaOH pH 7.5, 2 mM magnesium acetate, 0.1 mM TCEP, 0.1% v/v NP-40), supplemented with EDTA-free protease inhibitor tablets and 3 mg DNaseI. Cells were lysed by forcing the suspension through a needle with a 20 ml syringe 20 times. Cell lysate was cleared by centrifugation at 200,000 *g*, 4 °C for 25 min. The supernatant was filtered through a 0.6 PVDF filter and supplemented with 3 ml BioLock solution before binding in batch to 10 ml bed-volume equilibrated Strep-resin for 1 h at 5 °C. Beads were separated from the supernatant via a gravity flow column, washed one time with 20 ml lysis buffer and four times with 20 ml wash buffer (150 mM NaCl, 40 mM HEPES-NaOH pH 7.5, 2 mM magnesium acetate, 0.1 mM TCEP). Protein was eluted by adding five times 10 ml elution buffer (150 mM NaCl, 40 mM HEPES-NaOH pH 7.5, 2 mM magnesium acetate, 0.1 mM TCEP, 5 mM Desthiobiotin). All five fractions were pooled and diluted with 40 mM HEPES-NaOH pH 7.5 to a final NaCl concentration of 80 mM. The complex was then loaded on a 5-ml HiTrap Q HP column, equilibrated in buffer A (80 mM NaCl, 40 mM HEPES-NaOH pH 7.5, 2 mM magnesium acetate, 0.1 mM TCEP) and eluted with a 12 CV linear gradient to 100% buffer B (1 M NaCl, 40 mM HEPES-NaOH pH 7.5, 2 mM magnesium acetate, 0.1 mM TCEP). Fractions of interest were pooled, diluted with 40 mM HEPES-NaOH pH 7.5 to a final NaCl concentration of 80 mM and loaded on to a 1-ml ResourceQ column equilibrated in buffer A. The complex was eluted in a steep gradient (10 CV to 70% buffer B) to concentrate the complex. Selected fractions were pooled, flash frozen in liquid nitrogen and stored at −80°C until further usage.

#### *In vitro* transcription and poly(A) tailing

The PCR template for *in vitro* transcription contains the SP6 promotor sequence followed by the coding sequence of our transcript, encoding for an N-terminal 3x FLAG tag, a villin head piece domain and the cytosolic domain of Sec61b. T1 mix (1.3x stock; 40 mM HEPES-KOH pH 7.5, 6 mM MgCl_2_, 2 mM spermidine, 10 mM reduced glutathione, 0.5 mM ATP pH 7.3, 0.5 mM CTP pH 7.3, 0.5 mM GTP pH 7.3, 0.5 mM UTP pH 7.3, 0.1 mM CAP), PCR template (final concentration: 13 ng/μl), RNasin (final concentration: 1x) and SP6 polymerase (final concentration: 0.4 U/μl) were mixed and incubated for 90 min at 40 °C. RNA was purified with an RNAeasy kit (Qiagen) and subsequently poly(A)-tailed. For that 23 μg RNA (ca. 400ng/μl) was mixed with RNasin (final concentration: 1x), poly(A) polymerase (final concentration: 0.25 U/μl), ATP (final concentration: 1 mM) and 10x buffer (final concentration: 1x). The poly(A)-tailed, capped RNA was purified with a RNAeasy kit, flash frozen in liquid nitrogen and stored at −80°C until further usage.

#### *In vitro* translation

Homemade, endonuclease-treated, or -untreated rabbit reticulocyte lysate (RRL) was used as an *in vitro* translation system. The RRL was either supplemented with additional pig tRNA (0.1 mg/ml) (cT2) or no additional tRNA (cT2-tRNA) was added, to employ naturally occurring sub-optimal codons in *Oryctolagus cuniculus* for ribosome stalling on our transcript. The RRL was used as a 2x stock and was mixed with poly(A)-tailed mRNA (final concentration: 3 ng/μl), KOAc (final concentration: 10 mM), human CCR4-NOT (concentrations as specified for cryoEM, crosslinking mass spectrometry and Western Blot interaction studies), human CNOT4 (concentrations as specified for cryoEM, crosslinking mass spectrometry and Western Blot interaction studies), L-Methionine (final concentration: 40 μM) topped up with MilliQ water. The translation reactions were incubated for 30 min at 32 °C.

### CryoEM sample preparation

For cryoEM sample preparation, a total volume of 4 ml cT2-tRNA translation reaction containing poly(A)-tailed *rare* RNA (final concentration: 3 ng/μl), KOAc (final concentration: 10 mM), human CCR4-NOT (final concentration: 500 nM), human CNOT4 (final concentration: 1 μM), L-Methionine (final concentration: 40 μM) was incubated at 32 °C for 30 min. The sample was chilled on ice for 5 min and then 1 ml of translation reaction was loaded onto a 3 ml sucrose-cushion. The cushion consisted of 0.5 ml of 25% sucrose in 1x RNC buffer (50 mM HEPES-KOH pH 7.6, 100 mM Potassium acetate, 2 mM magnesium acetate) buffer, supplemented with 3 mM BS3, and 1.5 ml 15 % sucrose in 1x RNC buffer layered on top. The cushion was spun in a TLA 100.3 rotor for 1 h at 4 °C at 100,000 rpm. Subsequently, the supernatant was removed and each pellet resuspended in 25 μl 1x RNC buffer, supplemented with 3 mM BS3 and incubated on ice for 30 min. The reaction was quenched by the addition of 50 μl 1x RNC buffer with 40 mM Tris-HCl, pH 7.5. The resuspended sample of all four tubes were pooled and incubated with 60 μl equilibrated FLAG resin (50% solution). The sample was incubated with the resin for 1 h at 4 °C in a rotator. The beads were washed five times with 300 μl 1x RNC buffer. The sample was eluted with 30 μl elution buffer (0.5 mg/ml 3xFLAG peptide in 1xRNC buffer).

### CryoEM grid preparation and data collection

Affinity-purified elongating ribosomes were vitrified on UltrAuFoil R1.2/1.3 300-mesh grids (Quantifoil) coated with graphene oxide (GO). First, gold grids were washed with deionized water, dried and subsequently glow-discharged for 5 min with an Edwards glow discharger at 0.1 torr and 30 mA. 3 μl of a 0.2 mg/ml GO suspension (from a 2 mg/ml stock, diluted in deionized water (SIGMA) was applied onto the glow-discharged grids, incubated for 1 min, blotted away, washed 3x by dipping the grids into 20 μl deionized water drops followed by blotting (the top-side of the grid was washed twice and the bottom-side of the grid was washed once).

3 μl sample at a concentration of 160 nM (A260 of 8) was applied onto the dry GO-coated grids, blotted for 4 or 7 sec (dataset 1 and 3, respectively), −15 blot force, 0 sec wait at 100% humidity, 4 °C, Whatman 595 blotting paper, with a Vitrobot Mark IV and plunge frozen into liquid ethane. Grids were stored until data-collection in liquid nitrogen.

Two datasets were collected with a Gatan K3 camera on Titan Krios microscopes in counting mode using EPU software in faster acquisition mode (AFIS). The first dataset was collected on Krios4 at eBIC (Diamond) and yielded 10,009 micrographs (105,000x magnification, pixel size= 0.829 Å, total dose 47.5 e^-^/Å^2^, 39 frames, resulting in 1.2 e^-^/Å^2^ dose/frame). The second dataset was collected in-house on Krios3 and yielded 22,753 micrographs (105,000x magnification, pixel size= 0.86 Å, total dose 48 e^-^/Å^2^, 40 frames resulting in 1.2 e^-^/Å^2^ dose/frame).

### CryoEM data processing

Datasets were processed with RELION 3.1 (*35*). Raw movies were corrected with MotionCorr, followed by CTF correction using gctf. Particles were picked using a low-pass filtered 80S ribosomes as a 3D reference, resulting in 679,431 initial particles for optics group (og) 1 and 2,440,187 initial particles for og2 which were used for initial 2D classification. Good 2D classes were selected and subjected to 3D classification without alignment using data to 8.29 Å and 8.6 Å, respectively, which resulted in 325,837 high-resolution 80S particles for og1 and 2,181,772 80S particles for og2. Extracting the particles at full pixel size resulted in a 3D reconstruction at 2.8 and 2.9 Å, respectively.

To select for ribosomes translating *rare* mRNA and to exclude idle ribosomes, we performed focused classification with signal subtraction (FCwSS) on the P site, resulting in 48,653 (og1) and 253,945 (og2) P-site tRNA-containing ribosomes. Particles from og2 were subjected to another round of 2D classification, this time without alignment to remove noise and low-resolution particles. To identify the subset of elongating ribosomes bound to CCR4-NOT, we performed two consecutive FCwSS on the E site and selected 10,440 particles containing CNOT3 in og1 and 8,997 particles in og2, respectively. Particles were further refined using 3D refinement, particle polishing and CTF refinement. Both CNOT3-containing classes from og1 and og2 were combined and refined together, resulting in a 3.1 Å map from 19,437 particles. In parallel, we also re-extracted and refined and polished these particles again, but with a 2-fold downscaled pixel size of 1.658 Å/pix, which yielded a 3.3 Å map with improved map interpretability near the E site. This map (which has been deposited to the EMDB entry as an additional map) was used for model interpretation and figure making (Fig. 2B, S4A-B).

### Model building, refinement and validation

The molecular model from PDB 6SGC (80S stalled on a poly(A) stretch (*28*)) was split into three groups, which were individually docked into the 3.1 Å post-processed map using UCSF Chimera (version 1.15) (*36*). All chains were manually adjusted into the original, or suitably blurred maps (B factors of 50 to 200) using Coot (version 0.9.6, Marina Bay) (*37*). AlphaFold2 (*29, 30*) models for CNOT3 helical bundles *a* (residues 1-115) and *b* (116-236) were rigid-body docked in Coot and merged into group 2. Register errors were corrected manually based on the cryoEM density as was the linker connecting the two helical bundles. The nascent chain and P-site mRNA bases from PDB 6SGC were mutated to conform to the expected sequences. Owing to a lack of structures of tRNA^Leu,UUA^, we adjusted the sequence, anticodon stem loop and CCA end of the original tRNA^Lys,3^ from PDB 6SGC to fit a consensus sequence derived from *O. cuniculus* tRNA^Leu,UUA-1-1^, ^-2-1^ and ^-3-1^. Owing to the lower local resolution, the L1 stalk and protein uL1 were rigid body fitted into the cryoEM density.

In Phenix (version 1.20-4459-000) (*38*), the three groups were first combined using *iotbx.pdb.join_fragment_files* and *phenix.real_ space_ refine* was used to perform real space refinement of the resulting model with default settings and the following additions: (i) a parameter ‘.eff’ file was defined to connect the 3’ oxygen of A76 of the peptidyl tRNA to the carbonyl C of the C-terminal leucyl residue of the nascent chain via an ester bond geometric restraint of 1.4±0.025 Å; (ii)*phenix.elbow* was used to automatically obtain restraints for all non-standard RNA bases and ligands; (iii) a nonbonded weight of 1000 was used; (iv) rotamer outliers were fixed using the Fit option ‘outliers_or_poormap’ and the Target was set to ‘fix_outliers’; and finally, (v) 112 processors were used to speed up the calculations. A similar procedure was carried out for the 2-fold downscaled map.

### Molecular graphics

Map and model figures (Fig. 2, 3, 4, S2, S3, S4, S5) were generated using UCSF Chimera (version 1.15) (*36*), UCSF Chimera X (version 1.3) (*39*) and Pymol (version 2.4) (*40*), respectively. Graphics for all figures were created using Adobe Illustrator Creative Cloud 2022. Western immunoblots were cropped using Adobe Photoshop Creative Cloud 2022 and labeled in Illustrator. Unmodified blots are provided for review. Multiple sequence alignments (Fig. S5) were generated using Clustal Omega (*41, 42*) and jalview (version 2.11.2.3) (*43*). Crosslinks (Fig. S6) were illustrated using xiView (*44*). 2D class averages (Fig. S3) were generated in RELION 3.1 and the graph in Fig. S3C was plotted using GraphPad Prism (version 9.1.0, GraphPad Software Inc).

### Analytical sucrose density gradient centrifugation

300 μl *in vitro* translation reaction (50 nM CCR4-NOT and 75 nM CNOT4 variants final) were layered on top of a 12 ml 10-50% sucrose gradient and centrifuged for 2 h at 40,000 rpm, 4 °C, max acceleration, deceleration 9 in a SW 40 Ti swinging bucket rotor. Fractions were sampled with a Piston gradient fractionator and fraction profiles were generated by continuous absorbance measurement at 260 and 280 nm. Proteins in fraction 7-16 (45 μl, supplemented with 1x loading buffer), were analyzed on 12-well Bolt 4-12% Bis-Tris SDS-PAGE. Proteins were transferred onto a 0.2 μm nitrocellulose membrane with the BIO-RAD Trans-Blot Turbo Transfer System. Blots were blocked for 1 h in 5% milk in PBS at RT. First, blots were incubated with the rabbit primary antibodies (rabbit anti-CNOT4 1:500 (Abcam), rabbit anti-eS7 (Abcenta)) in 5% milk in PBS-Tween (PBS-T), overnight at 4 °C. After washing the membranes 3x with PBS-T, the blots were incubated with fluorescently labelled goat anti-rabbit antibody (1:5000, Invitrogen) for 1 h at RT and after washing with PBS-T. Blots were scanned with a LI-COR Imaging system at 600 and 800 nm. Second, blots were briefly washed with PBS-T and incubated with mouse anti-CNOT3 (1:5000, Santa Cruz Biotechnology) for 1 h at RT in 5% milk in PBS-T, washed 3x with PBS-T, incubated for 1 h at RT with fluorescently labelled goat anti-mouse (1:10000, Invitrogen) in 5% milk in PBS-T, washed and scanned as described above.

### Isotope labelled translation reactions

To monitor translation, ^35S^L-Methionine was added instead of cold L-Methionine to initiate the translation reaction. 19 μl radioactive translation reaction was supplemented with 1 μl ^35S^L-Methionine, incubated 30 min at 32 °C and layered on a 200 μl 10-50% sucrose gradient in RNC buffer. The gradient was spun at 55,000 rpm for 10 min at 4 °C in a TLS-55 rotor. Eleven 20 μl fractions were sampled and loaded on a 4-12% Bis-Tris NuPAGE and translated protein was detected *via* auto-radiography.

### Crosslinking mass spectrometry

For crosslinking mass spectrometry, 3 x 12 mL of translation reaction (6 x 2 ml) were analyzed. Each reaction contained 1 x cT2-tRNA lysate, 3 ng/μl *in vitro* transcribed, poly(A)-tailed *rare* mRNA, 10 mM Potassium acetate, 500 nM CCR4-NOT and 1 μM CNOT4. Addition of the proteins added ~40 mM NaCl. The reaction was started by the addition of 40.8 μM L-Methionine. The reactions were incubated for 30 min at 32 °C and kept on ice for 10 min before loading 2 ml of translation reaction on 10 ml 15% sucrose in 1 x RNC buffer each (50 mM HEPES-KOH pH 7.4, 100 mM Potassium acetate, 5 mM magnesium acetate). Ribosomes were pelleted for five hours at 38,000 rpm and 4 °C in a SW40 Ti swinging bucket rotor (max acceleration, deceleration 9).

After centrifugation, the sucrose was removed, and the pellet was resuspended in 150 μl 1 x RNC buffer on ice. For the first 12 ml, 4 ml of the reaction were crosslinked with 1 mM, 2 mM or 3 mM BS3, respectively for 30 min on ice. For the second and third 12 ml, 4 ml of the reaction were crosslinked with 0.5 mM, 1 mM or 2 mM BS3, respectively, for 30 min on ice. The crosslinking reactions were quenched by the addition of 5 mM Ammonium bicarbonate. The different crosslinking concentrations of the 12 ml were pooled (900 μl total) and incubated on 800 μl 50% modified Cytivia StrepTactin Sepharose High Performance resin. The resin was treated as previously published (*45*), to avoid Streptactin cleavage upon on-bead digestion of the sample. The sample was incubated for 45 min on ice, while the beads were agitated every 5 min to keep them in suspension. The beads were washed five times with 500 μl 1 x RNC buffer. The beads were subsequently kept on ice overnight.

The next day, the beads were re-suspended in 200 μl denaturation buffer (8 M Urea in 100 mM Ammonium bicarbonate (ABC)). The following steps, including trypsin digestions, were performed under constant agitation of 14,000 rpm in a heating block.

Reduction buffer (200 mM Dithiothreitol (DTT) in 50 mM ABC) was added to a final concentration of 10 mM DTT and incubated for 15 min at 25 °C. Subsequently, alkylation buffer (500 mM iodoacetamide (IAA) in 50 mM ABC) was added to a final concentration of 40 mM IAA and incubated for 30 min at 25 °C in the dark. 10 mM DTT was added to quench residual IAA and 0.5 μg LysC protease (MS grade, Pierce) was added for 4 hours at 25 °C. The sample was diluted with digestion buffer (100 mM ABC) to ensure a urea concentration below 1.5 M. 4 μg trypsin (MS grade, Pierce) was added to the sample and incubated for 16 hours at 16 °C. The digestion was stopped by the addition of 10% Trifluoroacetic acid (TFA) to pH 2.5. Peptides were cleaned-up via stage-tipping (*46*) and vacuum concentrator dried peptides were stored at −80 °C until further use.

100 μg of peptides were resuspended in size exclusion chromatography (SEC) buffer (30% acetonitrile (ACN), 0.1% TFA) and separated on a Superdex 30 Increase 3.2/300. The flow rate was 10 μl/min, and 50 μl fractions were collected after the void volume. Fractions containing crosslinked peptides (5-10) were dried in a vacuum concentrator and stored at −80 °C until further use.

### LC-MS/MS acquisition of crosslinked samples

Samples for analysis were resuspended in 0.1% v/v formic acid, 3.2% v/v acetonitrile. LC-MS/MS analysis was conducted in duplicate for SEC fractions, performed on an Orbitrap Fusion Lumos Tribrid mass spectrometer (Thermo Fisher Scientific, Germany) coupled online with an Ultimate 3000 RSLCnano system (Dionex, Thermo Fisher Scientific, Germany). The sample was separated and ionized by a 50 cm EASY-Spray column (Thermo Fisher Scientific). Mobile phase A consisted of 0.1% (v/v) formic acid and mobile phase B of 80% v/v acetonitrile with 0.1% v/v formic acid. Flowrate of 0.3 μl/min using gradients optimized for each chromatographic fraction from offline fractionation ranging from 2% mobile phase B to 45% mobile phase B over 122 min, followed by a linear increase to 55% and 95% mobile phase B in 2.5 min, respectively. The MS data were acquired in data-dependent mode using the top-speed setting with a 2.5 second cycle time. For every cycle, the full scan mass spectrum was recorded in the Orbitrap at a resolution of 120,000 in the range of 400 to 1,600 m/z. Normalized AGC = 400%, Maximum injection time = 50 ms, Dynamic exclusion = 60 s. For MS2, ions with a precursor charge state between 4+ and 7+ were selected with highest priority and 3+ were fragmented with any cycle time remaining. Normalized AGC target = 220%, Maximum injection time = 150 ms. Fragmentation was done with stepped-HCD collision energies 26, 28 and 30 % and spectra were recorded at 60,000 resolution with the orbitrap.

### Analysis of crosslinked MS data

A recalibration of the precursor m/z was conducted based on high-confidence (<1% FDR) linear peptide identifications. The recalibrated peak lists were searched against the sequences and the reversed sequences (as decoys) of crosslinked peptides using the Xi software suite (version 1.7.6.4) (*47*) (https://github.com/Rappsilber-Laboratory/XiSearch) for identification. The following parameters were applied for the search: MS1 accuracy = 2 ppm; MS2 accuracy = 5 ppm; Missing Mono-Isotopic peaks = 2; enzyme = trypsin (with full tryptic specificity) allowing up to three missed cleavages; crosslinker = BS3 (with an assumed reaction specificity for lysine, serine, threonine, tyrosine and protein N termini); Noncovalent interactions = True; Maximum number of modifications per peptide = 2; Fixed modifications = carbamidomethylation on cysteine; variable modifications = oxidation on methionine, hydrolyzed / aminolyzed BS3 from reaction with ammonia or water on a free crosslinker end. The database used was all proteins identified with an iBAQ > 1e6 (729 proteins) plus the translated nascent peptide.

Prior to FDR estimation matches were filtered for those with at least 4 matched fragments per peptide. The candidates were filtered to 2% FDR on residue pair level on link level using XiFDR version 2.2.beta.

## Acknowledgments

We are grateful to members of the Passmore lab, the Ramakrishnan lab and the Rappsilber lab for assistance and advice; Fabian Schildhauer for assistance with modified Streptavidin bead preparation; Ramanujan Hegde for the rare mRNA plasmid template; the MRC Laboratory of Molecular Biology Electron Microscopy Facility for access and support of electron microscopy sample preparation and data collection; J. Grimmett and T. Darling (LMB scientific computation); and J.G. Shi (baculovirus) for support. We acknowledge Diamond Light Source for access to eBIC (proposal BI23268) funded by the Wellcome Trust, MRC and Biotechnology and Biological Sciences Research Council. This work was supported by the Medical Research Council, as part of United Kingdom Research and Innovation, MRC file reference number MC_U105192715 (L.A.P.); a Postdoc fellowship of the Deutsche Forschungsgemeinschaft (DFG, German Research Foundation) to E.A. V.C. was supported by V. Ramakrishnan whose funding was from the MRC (MC_U105184332), the Wellcome Trust (WT096570), the Agouron Institute, and the Louis-Jeantet Foundation. The Wellcome Centre for Cell Biology is supported by core funding from the Wellcome Trust [203149].

## Supplementary Figures and Figure Legends

**Fig. S1.**
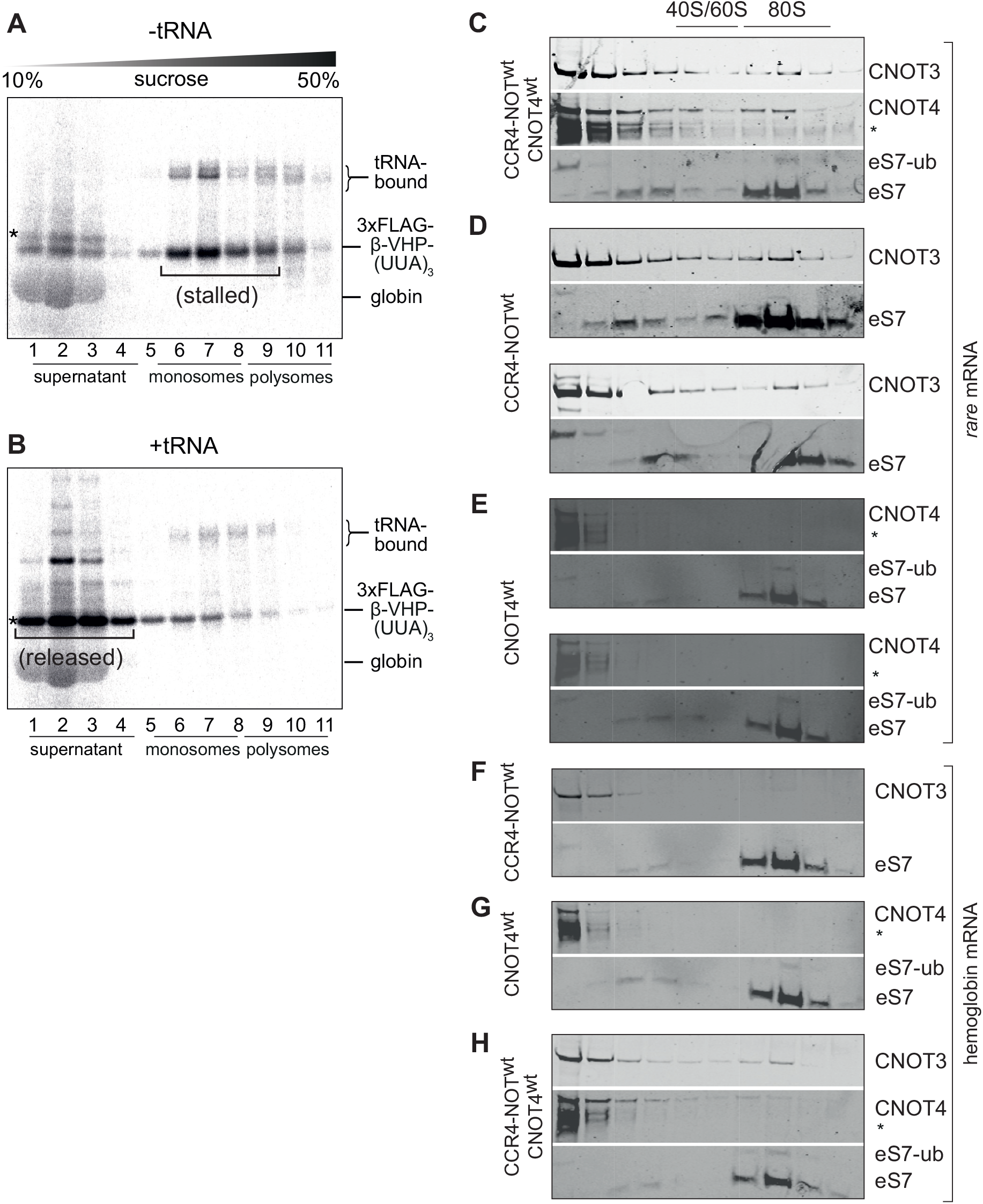
*In vitro* reconstitution of ribosome stalling on sub-optimal codons, CCR4-NOT/CNOT4 binding and eS7 mono-ubiquitination in the rabbit reticulocyte lysate. (**A**) Sucrose gradient fractionation of *rare* mRNA in RRL shows that the majority of the radiolabeled 3xFLAG-β-VHP-(UUA)3 is on stalled monosomes (fractions 5-8). * indicates a small amount of full-length protein product that was released from the ribosome at the stop codon and therefore migrated in the supernatant (fractions 1-3). We note that the previous cryoEM structure of yeast Ccr4-Not bound to RNCs stalled during elongation was obtained from the polysome fractions of a sucrose gradient. In our experiments, we observed mainly stalled monosomes. Multiple rounds of initiation are disfavored in the RRL owing to the high amounts of added mRNA and dilution of initiation factors during lysate preparation relative to intact cells. Consequently, only <5% of mRNAs have two or more ribosomes translating on a single mRNA (compare ^35S^L-methionine signals in monosome fractions 6-8 and polysome fractions 9-11). (**B**) The full-length protein (*) accumulates in the supernatant when *rare* mRNA is translated in RRL supplemented with total tRNA purified from pig liver. Autoradiograms in panels (A) and (B) show migration of the ^35S^L-Met-labelled nascent polypeptide. (**C-E**) Sucrose gradient fractions of non-radiolabeled translation reactions of *rare* mRNA in the presence of the indicated CCR4-NOT and/or CNOT4 proteins immunoblotted using antibodies against CNOT3, CNOT4 and eS7. The mono-ubiquitinated eS7 band is also labeled. (**F-H**) Same experiments as in panels C-E but performed on ribosomes elongating on α- and β-globin mRNA. Asterisks denote cross-reacting bands. We note that in previous studies in yeast, eS7 is mono-ubiquitinated in monosome fractions and poly-ubiquitinated in polysome fractions. Similarly, we observe eS7 mono-ubiquitination in the monosome peak but no poly-ubiquitination. We conclude that poly-ubiquitination is not required for stable binding of CCR4-NOT and CNOT4.

**Fig. S2.**
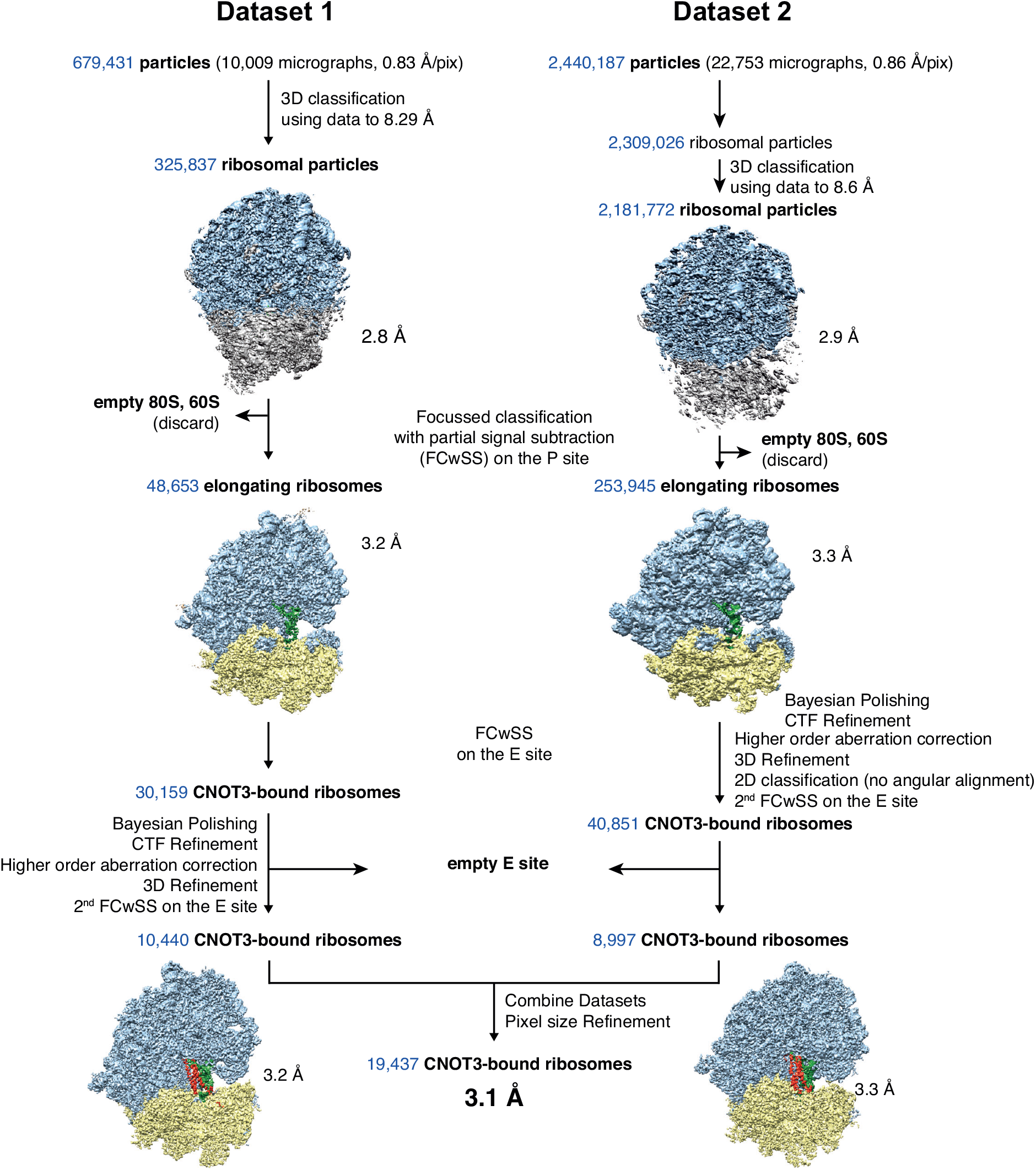
RELION 3.1 processing workflow. Two datasets were collected, processed separately and merged at the end to yield a final reconstruction at 3.1 Å resolution.

**Fig. S3.**
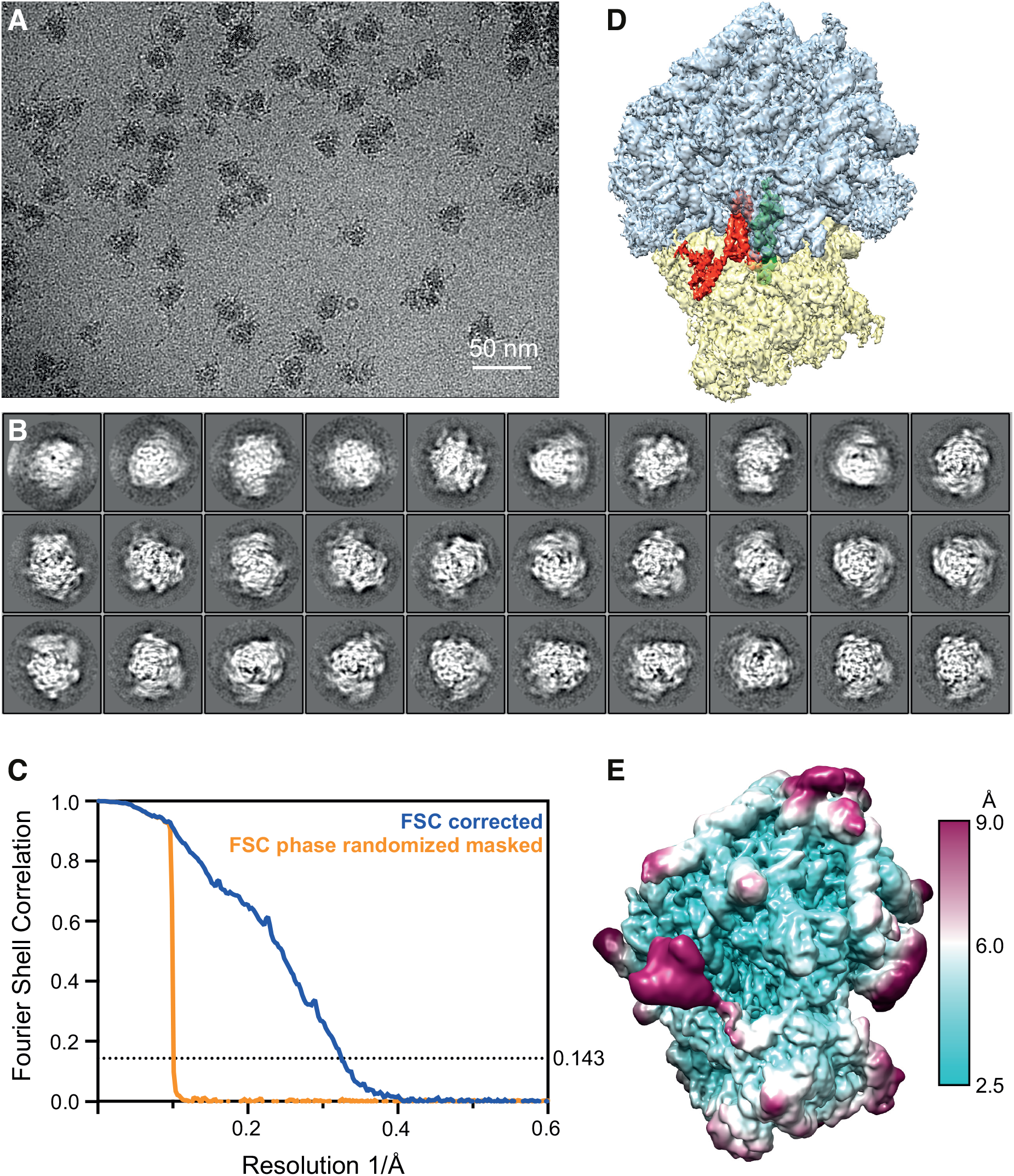
CryoEM analysis of stalled 80S ribosomes bound to CCR4-NOT and CNOT4. **A**) Representative micrograph used for single particle analysis. Scale bar: 50 nm. (**B**) Handpicked 2-dimensional class averages of 80S ribosomes (Table S1). (**C**) Gold-standard Fourier shell correlation (FSC) curve (blue) of the final map illustrating an overall resolution of 3.1 Å. The phase-randomized, masked FSC curve (orange) is also shown. (**D**) Final map segmented to show 60S (cyan), 40S (yellow), CNOT3 (orange) and the P-site tRNA (green). (**E**) Map from panel (D) colored according to the local resolution (cyan - high resolution; magenta - low resolution).

**Fig. S4.**
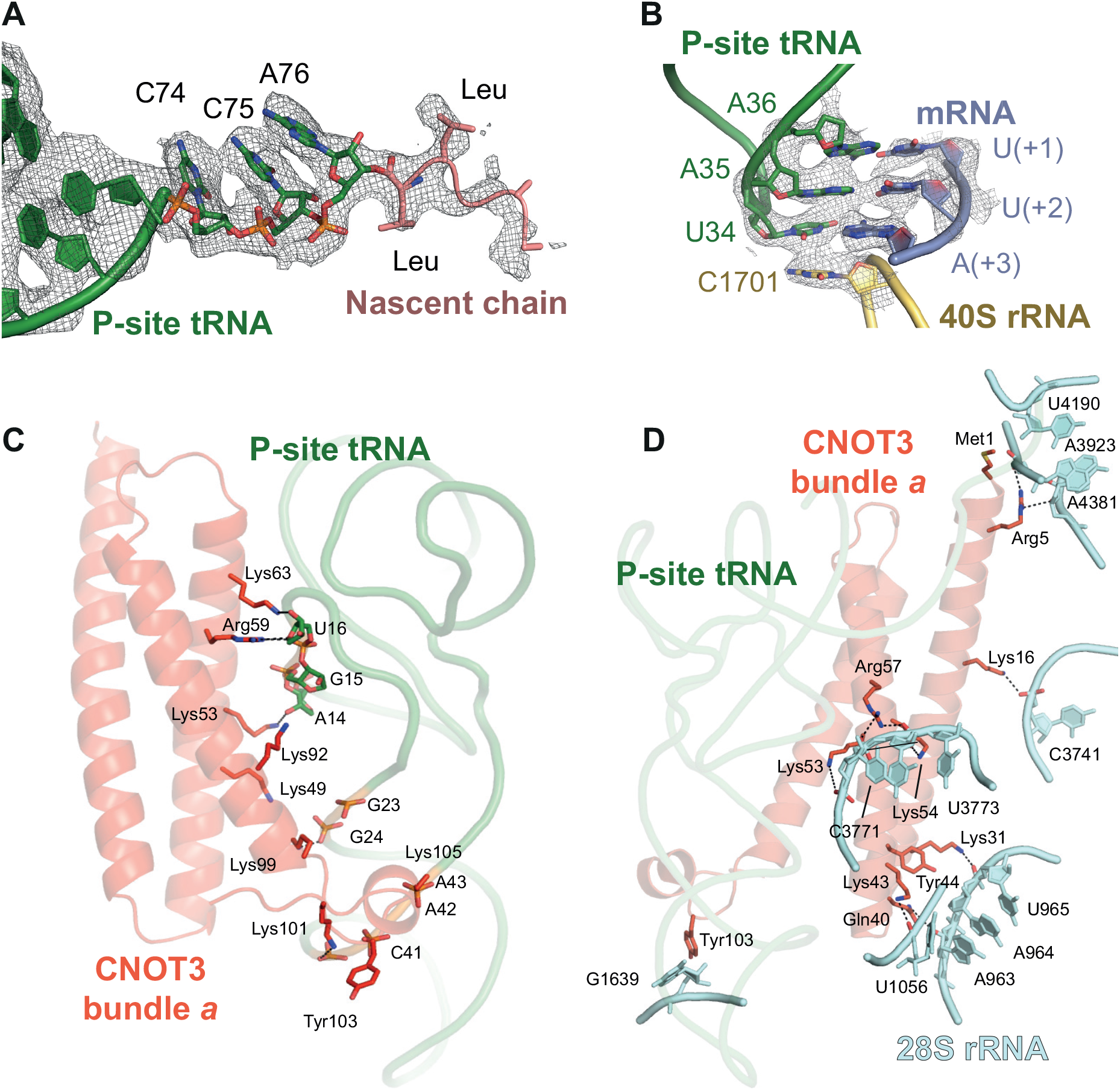
Details around the P site tRNA in the cryoEM map of 80S with CCR4-NOT. (**A-B**) The cryoEM map and atomic model are shown for the nascent chain (**A**) and the codon-anticodon in the P site (**B**). This map was calculated from 2-fold downscaled particles (to a pixel size of 1.66 Å/pix) to improve interpretability. The P-site tRNA and nascent chain are contoured at 10 r.m.s.d. (**C**) Packing interactions between CNOT3 helical bundle *a* (orange) and the P-site tRNA (green). (**D**) Packing interactions between CNOT3 helical bundle *a* (orange) and the 28S rRNA (cyan). Salt-bridges are indicated with black, dashed lines.

**Fig. S5.**
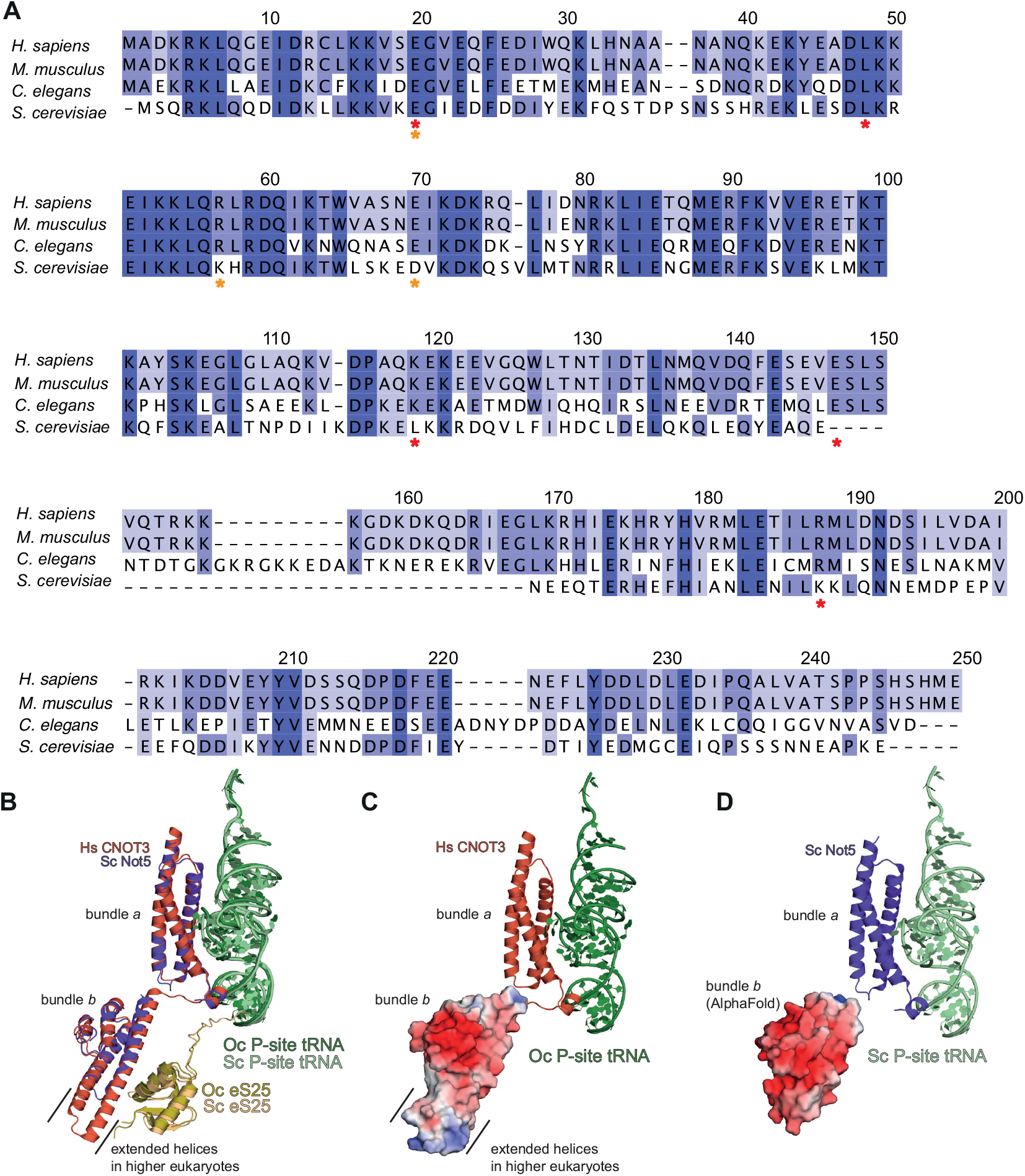
Evolutionary conservation of the interaction between CNOT3 and the 80S ribosome. (**A**) Multiple sequence alignments of the two helical bundles across representative eukaryotes (bundle *a*: 1-111, bundle *b*: 115-236). Numbering is according to the human sequence and frequently mutated residues in developmental disorders and cancer are indicated with red and orange asterisks, respectively. Helical bundle *a* has a sequence identity of 54.5% between yeast and human; helical bundle *b* (residues 120-213 in yeast Not5 and 115-236 in human CNOT3) has a sequence identity of 29.8%. High conservation, dark purple; lower conservation, lighter purple. (**B**) Superposition of budding yeast (Sc) Not5 helical bundle *a* (from PDB 6TB3) and helical bundle *b* (Alphafold2 model, residues 120-213) on human (Hs) CNOT3 helical bundles *a* and *b* (residues 1-236). Black lines indicate the extended helices in the human structure (residues 147-169), which are missing in yeast. Oc, *Oryctolagus cuniculus*. (**C**) Surface charge of helical bundle *b* of Hs CNOT3. (**D**) Surface charge of the AlphaFold2 prediction of helical bundle *b* of Sc Not5.

**Fig. S6.**
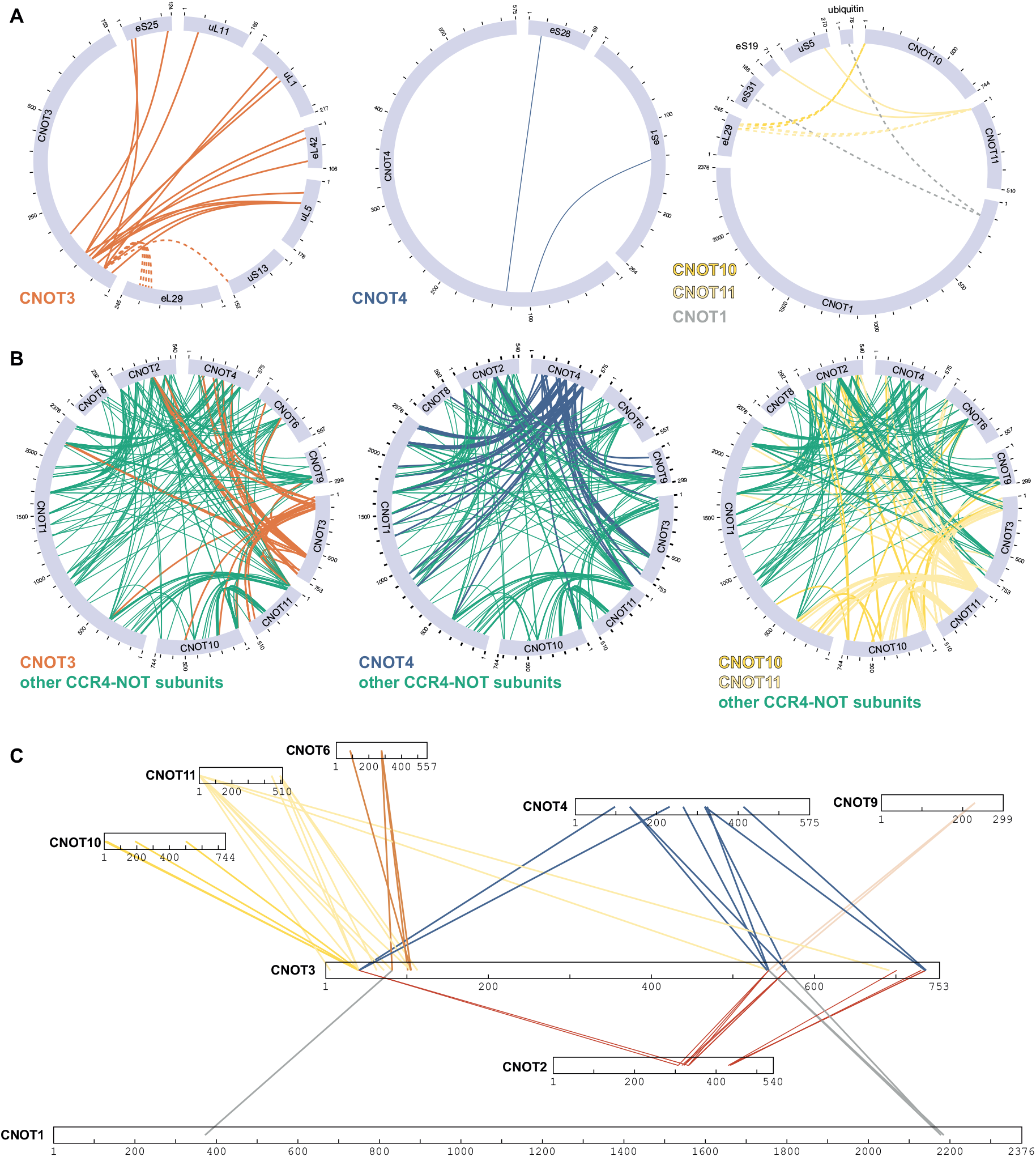
Crosslinking mass spectrometry of CCR4-NOT- and CNOT4-bound stalled 80S ribosomes. (**A, B**) Circle diagrams of observed crosslinks (**A**) between selected CCR4-NOT subunits or CNOT4 and 80S ribosomal proteins or (**B**) within CCR4-NOT/CNOT4. Dashed lines indicate crosslinks between proteins that were not visible in the structure and therefore could not be modelled on the structure. In panels (A) and (B), the indicated subunits are highlighted. (**C**) Diagrams of observed crosslinks of CNOT3 to other CCR4-NOT subunits and CNOT4, when bound to an 80S. CNOT3 is drawn to a larger scale than the other proteins.

**Fig. S7.**
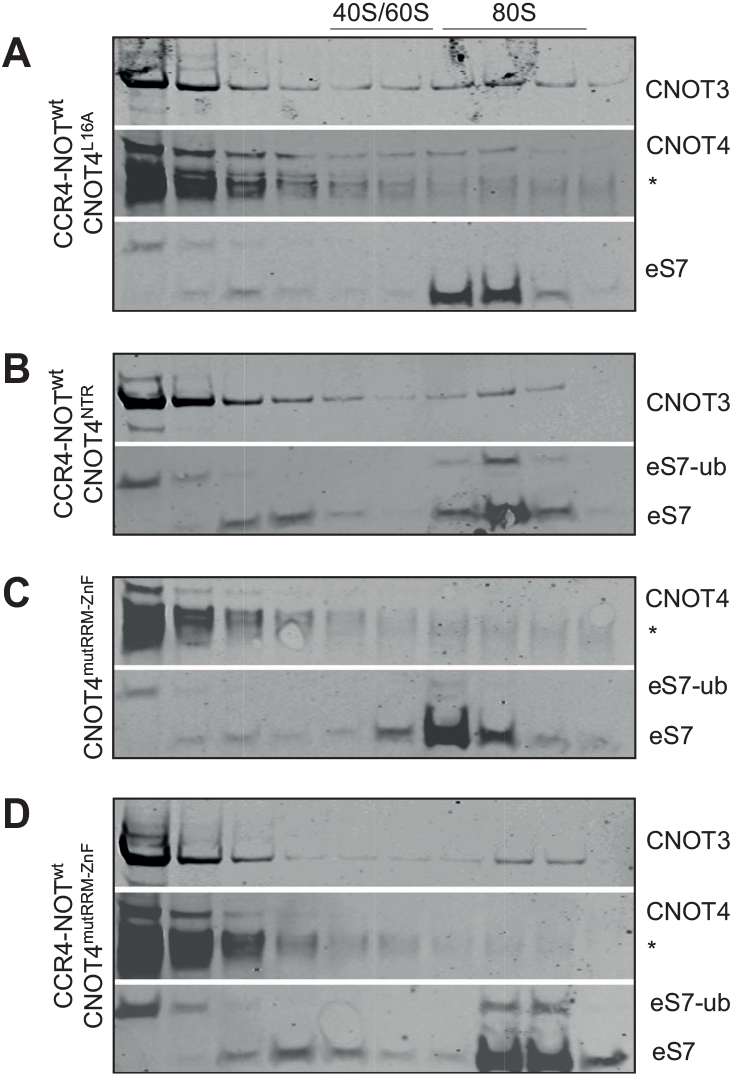
Repetitions of experiments in Figure 4

**Supplementary Table S1.** Data collection, processing, refinement and model statistics.

| Data Collection | Dataset 1 | Dataset 2 |
| --- | --- | --- |
| Voltage (keV) | 300 | 300 |
| Pixel size (Å) | 0.83 | 0.86 |
| Detector | Gatan K3 |  |
| Defocus range (µm) | -1.2 to -2.7 | -1.2 to -2.7 |
| Electron dose (e<sup>-</sup> frame<sup>-1</sup> Å<sup>-2</sup>) | 1.2 | 1.2 |
| Data Processing |  |  |
| Useable micrographs | 9,057 of 10,008 | 22,753 of 23647 |
| Particles picked | 679,431 | 2,440,187 |
| Final particles | 19,437 |  |
| Map sharpening B-factor (Å<sup>2</sup>) | -29 |  |
| Resolution (Å) | 3.1 |  |
| EMPIAR accession code | yyyyyy |  |
| EMDB accession code | EMD-xxxx |  |
| PDB accession code | aaaa |  |
| Model Composition |  |  |
| Chains | 84 |  |
| Non-hydrogen atoms | 219,018 |  |
| Protein residues | 12,159 |  |
| RNA bases | 5,649 |  |
| Metals (Mg<sup>2+</sup>/Zn<sup>2+</sup>) | 275/8 |  |
| Refinement |  |  |
| Resolution (Å) | 3.1 |  |
| CC (mask) | 0.73 |  |
| R.M.S deviations |  |  |
| Bond lengths (Å) | 0.009 |  |
| Bond angles (°) | 0.998 |  |
| Validation |  |  |
| Molprobity score | 1.56 |  |
| Clashscore, all atoms | 5.23 |  |
| Rotamers outliers (%) | 0.04 |  |
| Cβ outliers (%) | 0.01 |  |
| Ramachandran plot |  |  |
| Favored (%) | 95.91 |  |
| Allowed (%) | 4.04 |  |
| Outliers (%) | 0.05 |  |

